# Exhaustive benchmarking of *de novo* assembly methods for eukaryotic genomes

**DOI:** 10.1101/2023.04.18.537422

**Authors:** Dean Southwood, Rahul V Rane, Siu Fai Lee, John G Oakeshott, Shoba Ranganathan

## Abstract

The assembly of reference-quality, chromosome-resolution genomes for both model and novel eukaryotic organisms is an increasingly achievable task for single research teams. However, the overwhelming abundance of sequencing technologies, assembly algorithms, and post-assembly processing tools currently available means that there is no clear consensus on a best-practice computational protocol for eukaryotic *de novo* genome assembly. Here, we provide a comprehensive benchmark of 28 state-of-the-art assembly and polishing packages, in various combinations, when assembling two eukaryotic genomes using both next-generation (Illumina HiSeq) and third-generation (Oxford Nanopore and PacBio CLR) sequencing data, at both controlled and open levels of sequencing coverage. Recommendations are made for the most effective tools for each sequencing technology and the best performing combinations of methods, evaluated against common assessment metrics such as contiguity, computational performance, gene completeness, and reference reconstruction, across both organisms and across sequencing coverage depth.

## Introduction

Constructing a high quality, reliable genome assembly of an organism of interest enables a wide range of questions to be asked and answered. Annotation of gene regions enables for synteny analysis and comparative genomics, including posing evolutionary questions such as where and when genes have been gained or lost [1]. Chromosome-resolution assembly in particular allows for investigation of recombination, regulatory genomic architecture, and chromosome evolution [2]. It also enables effective and efficient use of emerging gene editing technologies such as the CRISPR/Cas-9 system, meaning a lack of, or low-quality, genome can hinder current and future manipulation, especially for species of economic and biological significance [3]. Undertaking a high-quality, whole genome assembly of a eukaryotic organism has become relatively affordable in the past 20 years, scaling from projects requiring millions of dollars and years of sequencing time such as that of the human genome [4 5], to a task which can be done on a budget of the order of a thousand dollars [6]. This exponential reduction in time and cost has fuelled a furious expansion of new sequencing methods and an explosion of algorithmic tools with which to process the resulting data.

However, despite the availability of these new sequencing technologies and a plethora of computational methods, there is no definitive guide available for researchers wishing to enter or navigate this space, particularly for those wishing to sequence the genomes of novel eukaryotic organisms. What combination of methods will produce the most biologically reliable, accurate genome, at minimal cost and with minimal computational resources? Is there a reliable workflow or schematic that can be adopted to bootstrap the process? What features of the genome of interest will determine what series of tools works best?

This is not necessarily a straightforward set of questions. Due to the increasingly widespread use of next-generation sequencing (NGS) technologies such as Illumina paired-end (PE) sequencing, as well as third-generation sequencing (TGS) technologies such as Oxford Nanopore MinION/PromethION, and PacBio continuous long reads (CLR) and circular consensus sequencing (CCS), laboratory teams have considerable options available by which to produce sequencing data. Coupled with a multitude of new genome assembly algorithms, determining what combination is best is difficult to predict *a priori*. It is unclear how particular algorithmic choices will affect the assembly of data from different technologies, and to what extent that assembly will depend on the genomic features of the organism being considered. Across these technologies, there are over 30 different algorithmic methods possible to produce a draft genome sequence of a eukaryotic organism, not including assemblies that could be obtained from merging drafts from other methods. Recent benchmarking studies vary, and understandably do not cover the whole field. Benchmark studies have typically focussed on certain organisms, such as prokaryotic genomes with long-read methods [7] or yeast with a variety of methods [8], or on certain technologies, such as Illumina methods [9 10], Nanopore methods [11 12], or PacBio methods [13 14]. Benchmarks are also available for other steps in the reference assembly pipeline, such as error correction of input reads across short reads [15] and long reads [16], as well as scaffolding draft genome assemblies using 3D chromosomal information through Hi-C sequencing [17]. The rapid development of novel algorithmic methods further complicates benchmarking, as new tools are not always benchmarked against the same combination of model organisms as other older methods, nor are they assessed against the same set of metrics. To the best of our knowledge, no benchmarking study has been performed comprehensively across Illumina, Nanopore, and PacBio technologies, as well as hybrid methods incorporating multiple of these, for eukaryotic genomes to date. Therefore, to fully understand the choices currently available to researchers, and to provide guidance on what methods should be chosen and when, the current market of possible methodologies needs comprehensive benchmarking.

We present an exhaustive benchmark of methods assembling two model eukaryotic genomes *de novo Caenorhabditis elegans* and *Drosophila melanogaster* using three current sequencing technologies: Illumina PE short reads, Nanopore long reads, and PacBio CLR sequencing. We include 20 different assembly packages, including both state-of-the-art and historical methods that have been used recently in the literature. In order to give a robust picture of how these methods perform, these assemblies are then polished using eight different polishing algorithms, in combinations based on sequencing technology input. At each stage, the assemblies produced by the methods are assessed across four types of metrics: contiguity and structural statistics, gene completeness, alignment to reference, and computational resource usage and performance, to provide additional guidance to researchers seeking to enter this area.

## Materials and Methods

### Rationale and scope

The construction of a high-quality reference assembly is a multi-step process; however, it is beyond the scope of any single benchmarking effort to comprehensively evaluate all tools across all steps. Therefore, we have limited the scope of the present benchmarking study to look at two key steps in current methods: the initial draft assembly of sequencing reads (short, long, or a hybrid of both), and the polishing of these draft assemblies using short reads, long reads, combinations of both methods in series, or hybrid methods using both concurrently. An overview of the approach taken is displayed in **Figure 1**. The methods selected for each step have been a combination of well-established methods routinely used in prior literature, and fresh methods developed recently which have as yet had limited literature presence. To hone the scope further, we have selected three prominent sequencing technologies to evaluate: Illumina PE short reads, Nanopore long reads, and PacBio CLR sequencing. These have been selected based on both their extensive use in recent assembly efforts, and the availability of public data for model organisms on which to benchmark. Newer technologies, such as PacBio CCS, also known as HiFi, as well as methods such as 10X Genomics sequencing, have been excluded due to limitations in sourcing sufficient public data.

**Figure 1:**
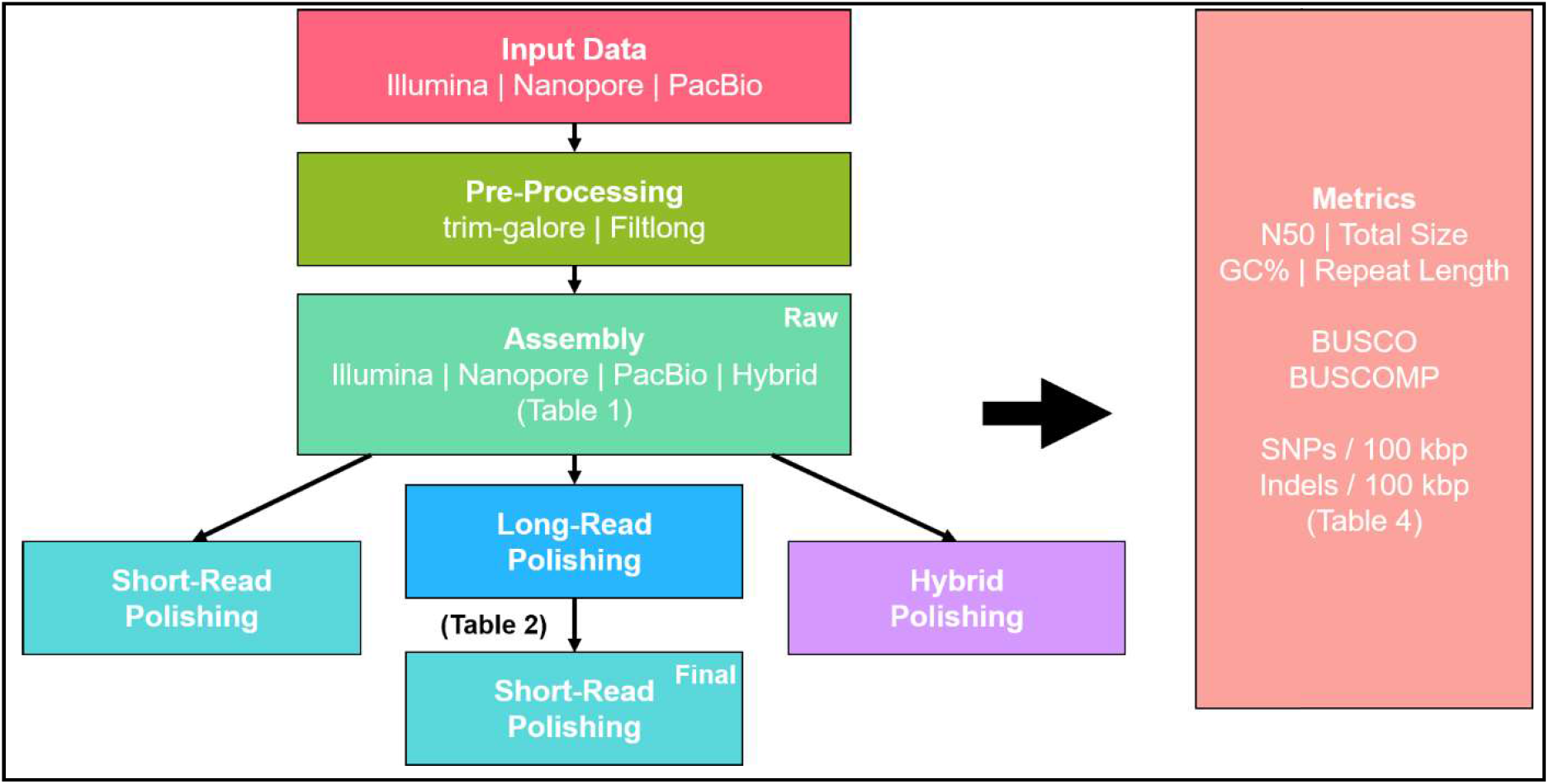
Flowchart of the rationale for the study. Data was collected from public repositories for Illumina, Nanopore, and PacBio sequencing technologies (**Table 3**). Illumina data was filtered and adapter-trimmed with trim-galore, while Nanopore and PacBio were trimmed and quality filtered with Filtlong. Processed reads were then assembled using each of the applicable assembly algorithms given in **Table 1**, producing a with Illumina assemblies only polished with short-read polishing (**Table 2**). Each assembly generated, raw or polished, was evaluated against several metrics (**Table 4**).

**Table 1:**
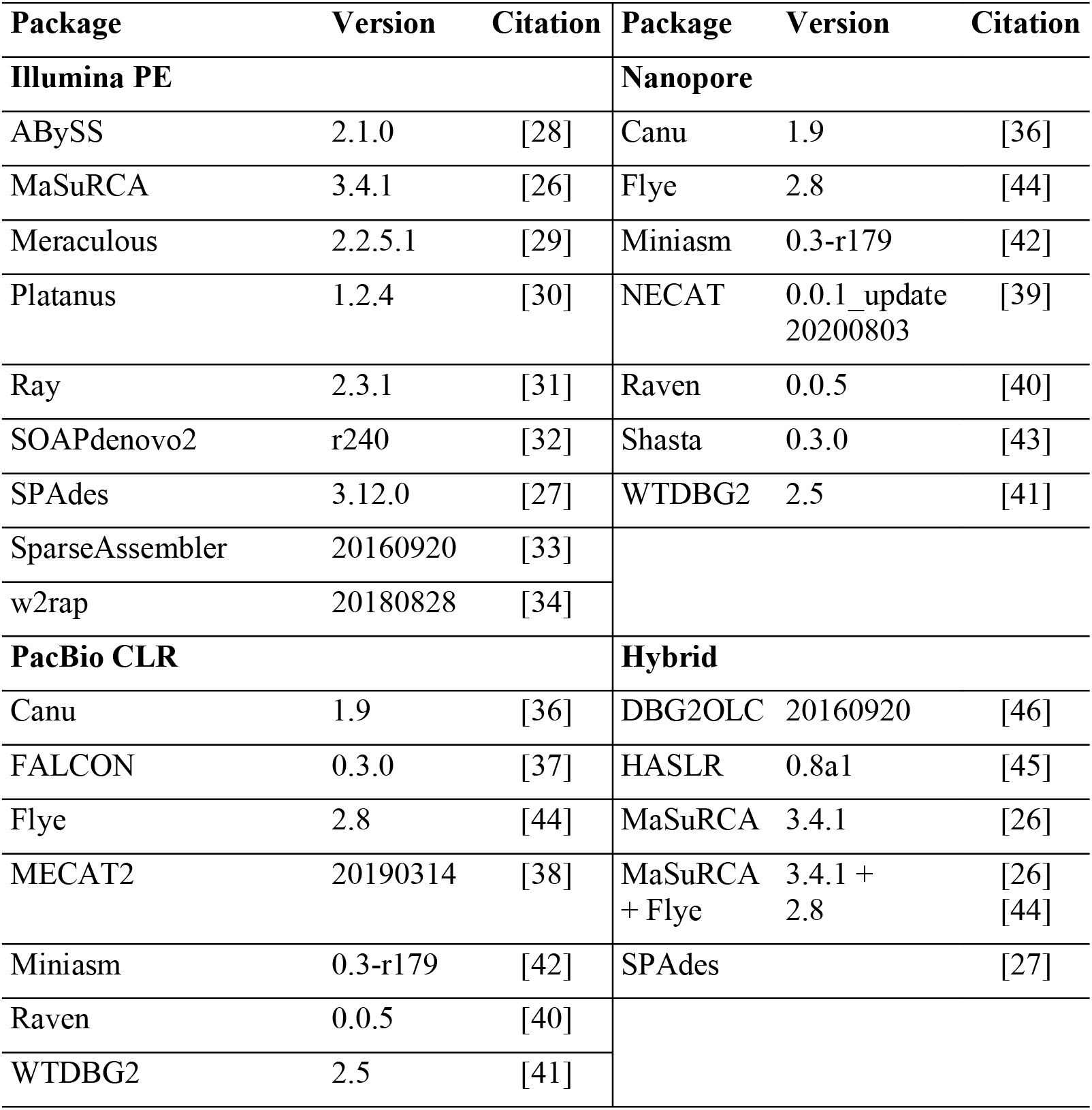
Assembly packages benchmarked in this study, grouped by accepted input data type

**Table 2:** Polishing packages benchmarked in this study, grouped by accepted input data type.

| Technology | Package | Version | Citation | Technology | Package | Version | Citation |
| --- | --- | --- | --- | --- | --- | --- | --- |
| **Illumina PE** | GATK | 4.1.3.0 | [50] | **Nanopore** | Medaka | 0.12.1 | [57] |
|  | HyPo | 1.0.3 | [51] |  | NextPolish | 1.3.0 | [52] |
|  | NextPolish | 1.3.0 | [52] |  | Racon | 1.4.13 | [56] |
|  | ntEdit | 1.3.2 | [53] | **PacBio CLR** | NextPolish | 1.3.0 | [52] |
|  | Pilon | 1.23 | [54] |  | Racon | 1.4.13 | [56] |
|  | POLCA | 3.4.1 | [55] | **Hybrid** | HyPo | 1.0.3 | [51] |
|  | Racon | 1.4.13 | [56] |  |  |  |  |

**Table 3:** Details of publicly available sequencing data used for benchmarking.

| Organism | Strain | Sequencing Technology | Source | Accession No. | Full Coverage |
| --- | --- | --- | --- | --- | --- |
| *C. elegans* | Wild | Illumina Paired-End | NCBI SRA | SRR9324028 | 48X |
|  | N2 | Oxford Nanopore | EBI ENA | ERR2092776, ERR2092777 | 80X |
|  | N2 | PacBio CLR | PacBio DevNet | - | 73X |
| *D. melanogaster* | ISO-1 | Illumina Paired-End | NCBI SRA | SRR6702604 | 41X |
|  | ISO-1 | Oxford Nanopore | NCBI SRA | SRR6821890, SRR6702603 | 57X |
|  | ISO-1 | PacBio CLR | PacBio DevNet | - | 98X |

**Table 4:**
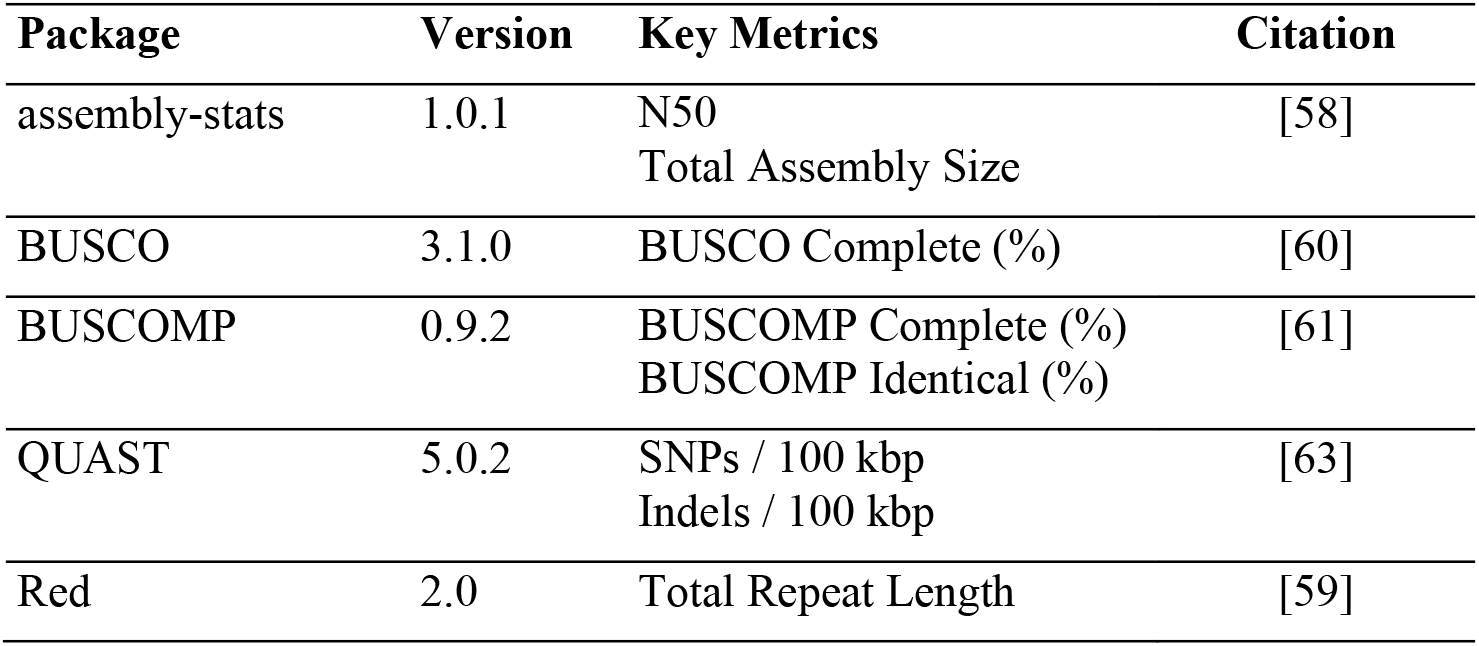
Metric-calculating packages used in this study, including the metrics extracted from each.

It is essential to benchmarking efforts to select materials in as unbiased a way as possible. Therefore, in this mindset, two organisms have been selected for benchmarking, both model eukaryotes used extensively in genetic and genomic studies in the literature: *Caenorhabditis elegans*, and *Drosophila melanogaster*. Both species have extremely high-quality, chromosome-level reference assemblies constructed from multiple sequencing technologies [18 19], including high-quality Sanger sequencing, data which has not been used in the answer is well established.

Based on these constraints, we have assembled *C. elegans* and *D. melanogaster* using 20 different assembly packages which, when accounting for packages supporting different input data independently or concurrently, provides 30 possible assembly methods overall (**Table 1**).

This is further broken down into nine methods for Illumina PE short reads, eight methods for Nanopore long reads, seven methods for PacBio CLR, and six hybrid methods which use two or more of the prior sequencing technologies concurrently during assembly. In addition, the assemblies produced by these algorithms have been polished with combinations of eight polishing packages which, when accounting for packages supporting different input data independently or concurrently, provides 12 possible initial methods overall (**Table 2**): seven methods using short reads, four methods using Nanopore long reads, two methods using PacBio CLR, and one hybrid method which uses short and long reads concurrently. These have been polished using multiple iterations of each polisher, to assess the benefit of repeated usage: long-read and hybrid polishing methods have been iterated four times, while short-read polishing methods have been iterated three times, extending upon common usage levels. In addition, to reflect current usage in the literature, all assemblies polished four times with long reads have been additionally polished with each short-read polishing method up to three times, to evaluate the effect of each combination of packages on the final quality of the assembly.

Two key interrelated components of assembly efforts are initial sequencing data collection and budget, which often interact and require compromise. Decisions must often be made *a priori* about the amount of sequencing data to collect which will provide a certain (often estimated) level of coverage of the genome of interest. We have therefore selected datasets which can be randomly subsampled to different levels of coverage, which can then be used to evaluate the extent of sequencing required to produce quality results within budget constraints (**Table 3**). The above combinations of assembly and polishing have been evaluated for three levels of coverage: two fixed levels of coverage 20X and 40X and one open level of coverage using full sets of data reasonably available, each providing more than 40X of coverage, with some data types containing up to almost 100X of coverage, which provides a realistic assessment of current high budget efforts.

Taking into account the above combinations, this study presents statistics for over 14,000 assemblies across *C. elegans* and *D. melanogaster*. Each assembly and its constituent methods have been assessed for quality across four categories of metrics (**Table 4**). First, we evaluated the contiguity and basic assembly statistics, including total assembly size, N50, repeat content, and GC content. Second, we assessed the gene completeness of the assembly, evaluated using benchmarked universal single-copy orthologues (BUSCOs). Third, we measured the accuracy of the assembly against the gold-standard reference for each organism, including the number of misassemblies, single-base mismatch errors, and insertion/deletion errors. Finally, we evaluated the computational performance of each package, in terms of memory requirements and total wall clock time.

To provide a fair assessment based on resources commonly available to research groups, all packages and methods have been run using a single 20-core computing node running Ubuntu 20.04, configured with 128 GB of RAM. To generate the levels of coverage for testing, the full data set for each technology for each organism was randomly subsampled using the BBMap reformat.sh package [20], estimating the desired total read length using genome sizes of 100 Mbp and 140 Mbp for *C. elegans* and *D. melanogaster*, respectively. The input data have been initially processed for basic quality and adapters using trim-galore [21] for Illumina data, and Filtlong [22] for long-read data, with details of parameters provided in the **Supplementary Material**. Each method has then been run on this processed data as close as possible to default or developer-recommended settings, except in rare cases where settings were adjusted with limited trial and error in order to produce an assembly. All assemblies were evaluated against the *WBcel235* reference for *C. elegans*, and the *r6* reference for *D. melanogaster*. The commands used for each package are provided in the **Supplementary Material**, with a brief overview of packages included in the study given below.

### Assembly algorithms

Two primary classes of algorithms exist for assembly, across short reads, long reads and hybrid algorithms, namely those based on de Bruijn graphs (DBG), and those based on an overlap-layout-consensus (OLC) approach, with some variation on both types of approaches unique to each package [23]. DBG-based approaches rely on constructing directed graphs, where *k*-mers generated from input reads form the edges, and *k*-mer overlaps form the nodes. As these approaches construct a graph from *k*-mers rather than whole reads, they are more prone to errors from input data, and struggle to resolve repeat regions when the *k*-mer size is small [24]. However, they are fast, scaling better with input data than OLC approaches [23], which is an important consideration given high coverage often obtained from sequencing. OLC approaches rely on constructing a graph of overlaps between reads, requiring a time-consuming overlapping step prior to reducing and resolving the graph [25]. However, these approaches are often more suited to handling errors, and provide more support over repetitive regions than DBG-based approaches, particularly for long reads [23]. Due to their high per-base quality and short length, short-read approaches tend to use a DBG-based approach to assembly. All nine algorithms considered here which rely solely on short-read input incorporate DBG-based approaches at some point in their algorithm (**Table 1**).

The primary difference between them is in whether reads are error-corrected before assembly, a step which takes additional time but can produce more contiguous assemblies, or whether a coverage cut-off is used instead to determine putatively erroneous reads. Of the nine assemblers, MaSuRCA [26] and SPAdes [27] rely on an error-correction step, while ABySS 2.0 [28], Meraculous [29], Platanus [30], Ray [31], SOAPdenovo2 [32], SparseAssembler [33], and w2rap [34] rely on a coverage cut-off.

Long-read assemblers have transitioned back towards OLC approaches, with some adjustments, in order to maximise the usage of longer read lengths while also better handling higher error profiles of reads [35]. Some long-read assemblers use an overlap-based error-correction step prior to assembly, in order to improve accuracy and, ideally, generate longer, more reliable contigs, which are then corrected again post-assembly using a consensus step. Others forgo the initial error-correction, while still maintaining the final consensus call. Others still forgo both error correction and consensus, providing draft genome assemblies at high speeds, but with the potential for higher errors. Of the nine long-read assemblers evaluated here, Canu [36], FALCON [37], MECAT2 [38], and NECAT [39] use error-correction steps before assembly along with consensus steps post-assembly. Raven [40] and WTDBG2 [41] forgo the error correction to speed-up assembly, but maintain a light-weight consensus step post-assembly. Miniasm [42] and Shasta [43] forgo both error correction and final consensus calls to produce rapid assembly. Flye [44] uses a unique approach relying containing a final consensus step.

In this study, we also evaluated hybrid methods which take both short-read and long-read input to construct the draft assembly. These methods often use a combination of DBG and OLC methods to construct the assembly, indicative of the multiple input types of data. The primary difference between the four packages considered here is in the use of short-read data within the assembly process [45]. One approach is to use the short-read data to correct the long-read input, before assembling the corrected long reads this is employed by MaSuRCA [26] in its hybrid assembly approach. Alternatively, the short reads can be assembled by a short-read assembler first, before aligning long reads to this graph to resolve ambiguities and generate longer contigs this is the approach taken by both SPAdes [27] and HASLR [45]. Finally, short reads can be assembled using a short-read assembler as in SPAdes and HASLR, but the resultant assembly graph is then used to compress the long reads before assembling them using an OLC-based approach this is the approach taken by DBG2OLC [46].

The assembly packages considered here have different requirements for input data, with some long-read assemblers designed for PacBio CLR data only, such as FALCON and MECAT2, or for Oxford Nanopore data only, such as NECAT and Shasta. The requirements and options used in this study are provided in **Table 1**, with further details available in the **Supplementary Material**. It is worth noting that assembly packages which require input data not used in this study, including packages designed specifically for PacBio HiFi reads, such as Peregrine [47] and HiCanu [48], or for 10X Genomics reads, such as Supernova [49], have been excluded from consideration.

### Polishing packages

Three types of polishing packages are considered here, categorised based on the input data they use in a particular instance (**Table 2**). There are seven packages which accept short-read-only input, namely the GATK pipeline [50], HyPo [51], NextPolish [52], ntEdit [53], Pilon [54], POLCA [55], and Racon [56]. In addition, two of these packages will accept long-read-only input as well: NextPolish and Racon, and have been considered in each category separately. A third long-read polishing algorithm, Medaka [57], is also considered for assemblies using Nanopore data. One package, HyPo, also explicitly accepts both short- and long-read input concurrently when polishing, and has been considered here as a hybrid polishing approach in addition to a short-read polishing package.

Polishing methods have been trialled in different iterations on the ‘raw’ assembly, specifically up to three iterations for short-read inputs, and four iterations for long-read inputs. In addition, short-read polishing methods have been trialled in combination on top of the (four times) long-read polished assemblies (**Figure 1**), reflecting common practice in recent literature. Details of each package are provided in the **Supplementary Material**.

### Evaluation metrics

Each assembly generated has been evaluated across four categories of metrics, with particular metrics of interest within each category calculated using various tools (**Table 4**). First, assemblies have been evaluated for contiguity and basic assembly statistics, such as N50, GC%, total repeat length, and total assembly size, using the assembly-stats [58] and Red [59] packages. These metrics provide an indication of how much material has been assembled and whether the material matches broad expectations based on the reference assembly. However, these statistics are calculated independent from the reference, and are useful for *de novo* efforts where no reference is available.

Second, the gene content has been indirectly assessed by analysing BUSCOs present in the assemblies, using both the BUSCO package [60], which analyses a single assembly using stringent criteria, and the BUSCOMP package [61], which takes BUSCO results for multiple assemblies and compares them based on alignments, with a more detailed breakdown of their contents. These have been calculated according to the OrthoDB v 9 [62] Nematoda and Diptera datasets, for *C. elegans* and *D. melanogaster* respectively.

Third, the gold-standard reference assembly has been used to assess the accuracy of the assembly and its potential errors. This is split into certain types of errors: number of misassemblies, number of mismatches or single-nucleotide polymorphisms (SNPs), and number of insertions/deletions (indels), calculated using QUAST [63]. These have their own unique impacts upon the quality of the assembly. Misassemblies can impact upon the quality of the scaffolded assembly when initial assembly and polishing methods are combined with a scaffolding package [64]. SNPs can affect upon per-base accuracy, impacting upon analyses such as evolutionary comparative genomics where single base changes are treated as putative mutations rather than assembly errors [65]. Indels can impact upon gene annotation efforts, where frame shifts can substantially change the translated content of the genome, or prevent detection by annotation algorithms [66].

Finally, assembly and polishing algorithms are assessed for their computational performance, explicitly broken down into total wall clock time and peak RAM consumption. While computing resources are becoming readily available and increasingly affordable, it is worthwhile knowing beforehand what scale of resources need to be requested, and for what period of time, and may prove a definitive metric for making decisions between two equally performing assembly or polishing packages.

In order to provide an overall ranking for the assemblies, z-scores are calculated for each primary metric across the pool of assemblies within each coverage level, for each organism, as is consistent with previous benchmarking efforts [10, 13]. In this study, the metrics included in the cumulative z-score calculation are namely: N50, where higher is ranked better; percentage of complete BUSCO genes, where higher is ranked better; percentage of identical BUSCOMP genes, where higher is ranked better; number of SNP errors per 100 kbp, where lower is ranked better; and number of indel errors per 100 kbp, where lower is ranked better. These are then summed and ranked within each coverage level for each organism, with: **Z**_cumul_.= **Z**_N50_ + **Z**_BU*c*_ + **Z**_BCP*id*_ − **Z**_SNP_ − **Z**_indel_.

## Results

The data generated in this benchmarking study is multi-dimensional, allowing for numerous interrogatory questions to be asked and investigated. Here we present our key findings from analysis of the data, broken down by organism: first, those for *C. elegans*, and second, those for *D. melanogaster*. For each organism, we present analysis of all three levels of coverage (20X, 40X, and full), and we explore three primary axes of the data. First, we present the performance and quality of assembly for each assembly package with no polishing, that is, raw assembly. Second, we present analysis of the effects of iterations of polishing upon assembly quality and accuracy across polishing packages using different input data types. Third, we evaluate the quality of so-called final assemblies for each combination of methods, to determine the extent to which each assembly algorithm can be improved by polishing, and what combination of methods works best for each.

### The *C. elegans* data set

*C. elegans* is a nematode which has been extensively used as a model organism, primarily to investigate neuronal development [67]. Its genome was the first whole-genome sequencing study for a multicellular organism [68] and has been extremely well characterised with extensive genomic resources available [18]. The diploid reference genome assembly used in this study, *WBcel235* [68], is approximately 100 Mbp in size, distributed across five autosomes, one heterochromosome, and a mitochondrial genome, and has a GC content of 35.4%, with a repeat content of 42.6% as measured by Red [59].

#### Contiguity and structural statistics

Contiguity did not change substantially between raw assemblies and polished ones, indicating that, by and large, the contiguity of the assembly is dictated by the assembly package used (**Figure 2**). Specifically, long-read assemblers produced the most contiguous assemblies, with consistently higher N50 values than short-read and hybrid methods (**Figure 2C D**). In addition, for long-read assemblies, contiguity tended to increase with coverage, though with diminishing returns in most cases. Hybrid methods performed well comparatively on low levels of coverage (20X), but were often overtaken by long-read methods for higher coverage. One particular assembler of note here is Flye, which produced highly contiguous assemblies at all coverage levels, for both Nanopore and PacBio input data, producing the highest contiguity assembly overall (N50 of 6.5 Mbp). The size of the assembly generated also did not increase with polishing, with some assemblies among the hybrid methods reducing in size with polishing, though not to a substantial extent (**Figure 2A B**).

**Figure 2:**
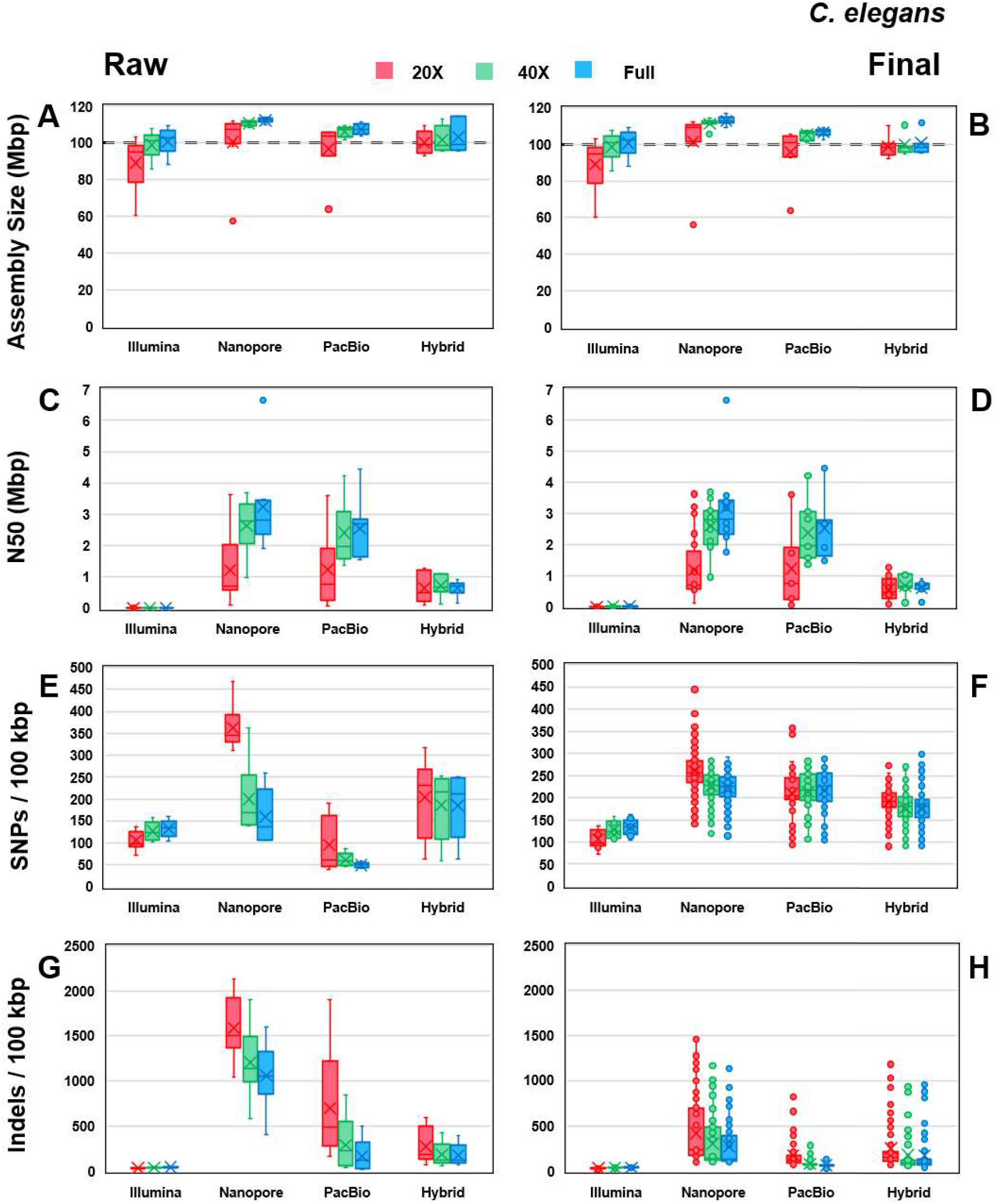
Box-and-whisker comparative plots for *C. elegans* for raw assemblies, i.e. without polishing, and final assemblies, i.e. polished with four rounds of long reads, followed by three rounds of short reads. Illumina assemblies are unpolished in both ‘raw’ and ‘final’ and are present for comparative purposes. The median is indicated by a horizontal line in each box, and the mean is indicated by a cross. Boxes represent the interquartile range of input assemblies. **Row 1:** Total assembly size for **(A)** raw and **(B)** final assemblies. **Row 2:** N50 for **(C)** raw and **(D)** final assemblies. **Row 3:** Average mismatch errors (SNPs) per 100 kbp of **(E)** raw and **(F)** final assembly sequence. **Row 4:** Average insertions and deletions per 100 kbp of **(G)** raw and **(H)** final assembly sequence.

Overall, the amount of reference genome covered by the assembly appears to be dictated by assembly algorithm, rather than polishing algorithm. Long-read assemblers tended to over-assemble at this stage, producing longer assemblies than the reference. Hybrid and Illumina methods tended to produce assemblies much closer to the expected genome size. However, assembly sizes for all input types tended to increase with additional coverage of data. This effect was particularly pronounced for particular assembly packages, such as Shasta and FALCON, which were not able to assemble more than 60-70% of the genome at 20X coverage, but produced more complete assemblies at 40X and full levels, indicating that these assembly algorithms require higher levels of support for individual regions of the genome before producing a consensus sequence.

The GC% of the assemblies did not change substantially with polishing in general, polishing produced tighter consensus between assemblers of the same dataset, indicating potential differences in the raw sequencing data itself, particularly between Nanopore and PacBio (**Supplementary Figure S1**). In this case, Nanopore produced assemblies with substantially higher GC% than other methods, while hybrid methods produced assemblies tightly clustered around the reference GC% level. There is some slight increase in GC% with coverage, but not to a large degree. Raw long-read assemblies tended to overestimate the total repeat length compared to the reference, with hybrid and Illumina methods producing results more in line with the reference. However, this was normalised somewhat by polishing, which reduced the length of repeat regions for both Nanopore and PacBio data.

#### Gene completeness

Hybrid assemblies tended to produce the most initially complete assemblies, but long-read assemblies supplemented with short reads at the polishing stage tended to overtake them in completeness (**Figure 3**). The presence of complete BUSCO genes was more readily identifiable in Illumina and hybrid assemblies, exhibiting values at the raw assembly stage that were similar to the final assembly stage (**Figure 3E**). However, for PacBio and Nanopore assemblies, polishing made a substantial difference in the identification of BUSCOs in an assembly, with this effect being more pronounced for low coverage levels. Illumina assemblies in this case peaked below 90% completeness measured by BUSCO, while assemblies incorporating longer reads managed to produce assemblies with over 99% completeness by the same metric once polished, despite initially lower numbers at the raw assembly stage. These results suggest that BUSCO complete scores provide a good indication of the effectiveness of polishing for these assemblies, and the progress towards the final assembly quality.

**Figure 3:**
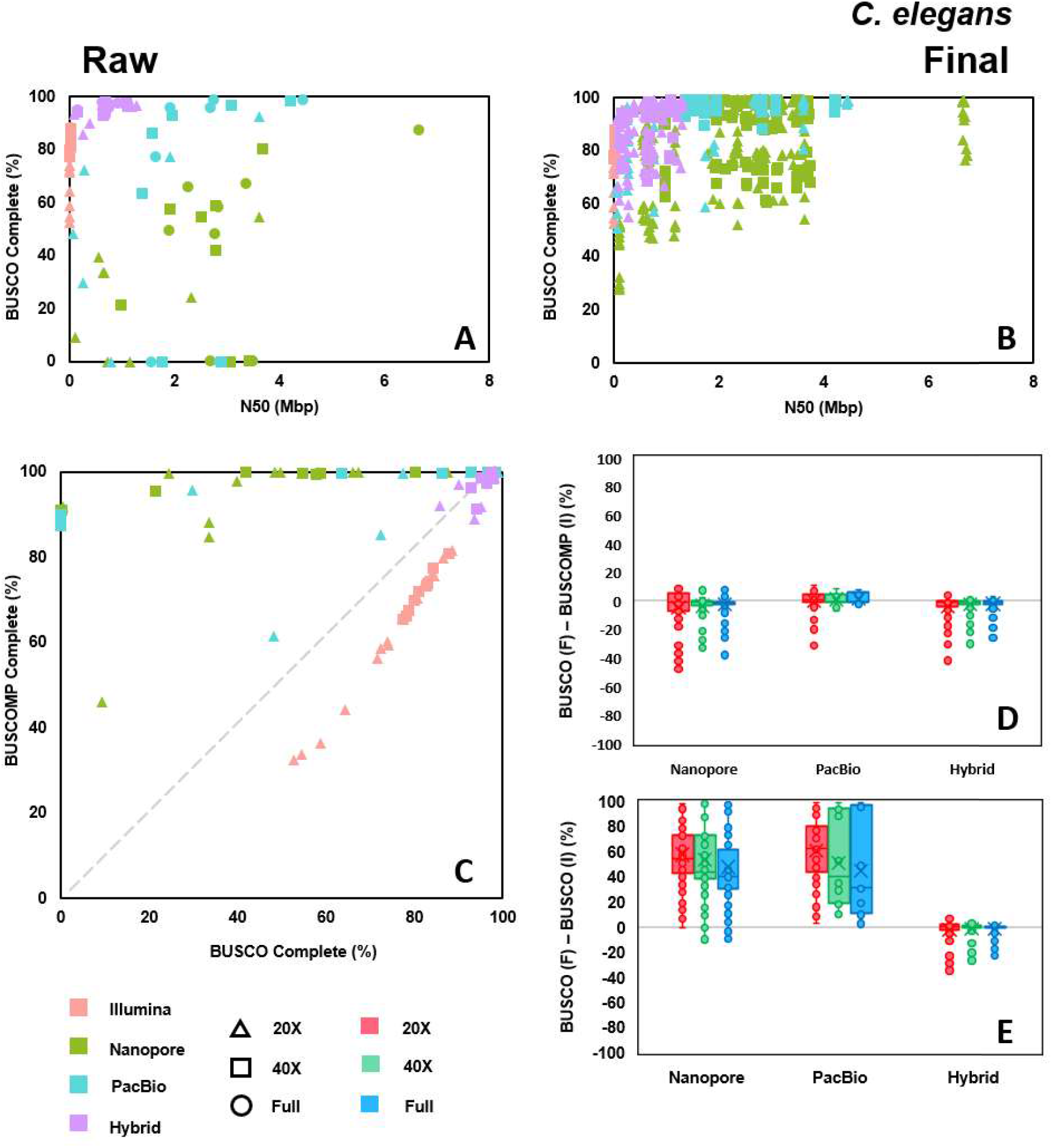
Comparison of complete BUSCO genes for C. elegans for raw assemblies, i.e. without polishing, and final assemblies, i.e. polished with four rounds of long reads, followed by three rounds of short reads. lllumina assemblies are unpolished in both ‘raw’ and ‘final’ and are present for comparative purposes. For the box-and-whisker plots, the median is indicated by a horizontal line in each box, and the mean is indicated by a cross. Boxes represent the interquartile range of input assemblies. **Top:** Percentage of complete BUSCO genes found in **(A)** raw and **(B)** final assemblies. **Bottom Left: (C)** Comparison of the percentage of complete BUSCOMP genes found in the raw assemblies against the percentage of BUSCO genes; the diagonal indicates equality between the two metrics. **Bottom Right:** Box-and-whisker plot comparisons by sequencing type of the difference between the percentage of **(D)** complete BUSCOMP genes or **(E)** complete BUSCO genes in the raw assembly, and the BUSCO genes in the final assembly, with the horizontal grey line indicating no change between raw and final. Negative values indicate overestimation by the raw assembly metric of the final assembly metric, while positive values indicate underestimation.

Results from BUSCOMP analysis tell a slightly different story. BUSCOMP completeness is substantially higher for long-read assemblies, likely due to an increased tolerance of errors when detecting the presence of BUSCOs compared to the BUSCO package. Of particular note is the difference between final BUSCO scores and initial BUSCOMP scores, and the potential to use them as a predictor of final quality; put another way, is the initial BUSCOMP completion score useful in predicting quality post-polishing? There tends to be a significantly lower difference between initial BUSCOMP score and final BUSCO score for long-read assemblies compared to differences between initial BUSCO score and final BUSCO score (**Figure 3D E**), indicating that the difference between these scores at the raw assembly stage is a useful measure of the potential to be gained from polishing that assembly.

Coverage made a substantial difference in the identification of complete BUSCOs in both raw and final assemblies (**Supplementary Figures S2 3**). However, this was less pronounced for a few assemblers, namely hybrid assembly methods along with SPAdes and SOAPdenovo2 for Illumina data, and Flye for PacBio data, which had less gain with additional coverage. Additionally, highly complete assemblies could still be produced even with low coverage, depending on the methods selected (**Supplementary Figures S4 5**). Notable high performers in this case are Flye, Raven, and WTDBG2 for Nanopore and PacBio data, and DBG2OLC and MaSuRCA+Flye for hybrid data, all of which were able to produce assemblies with over 98% of complete BUSCOs found with only 20X of coverage.

#### Accuracy

The most useful indicators of accuracy when comparing between potential assemblies were SNP and indel errors per 100 kbp, which seemed to be more robust to low overall assembly length and fragmentation, while metrics such as total number of misassemblies were not as robust. SNP errors showed a clear difference between Nanopore and PacBio assemblies, with Nanopore raw assemblies having significantly higher SNP errors than PacBio (**Figure 2E F**). Hybrid and Illumina methods had fewer SNP errors than Nanopore, but still higher than PacBio. Compared to the raw assemblies, polishing appears to increase SNP errors in some cases, particularly for higher coverage levels, and particularly for PacBio data. The exception to this trend is for low coverage Nanopore assemblies, where polishing tended to correct SNP errors. The short-read polisher used had a pronounced effect on final SNP rates, with ntEdit producing assemblies with the lowest SNP errors, followed by POLCA (**Supplementary Figure S7**). For POLCA in particular, coverage helped correct more SNP errors no matter the initial sequencing technology used for the assembly, a trend not seen across the majority of other polishing algorithms, where increased coverage tended to result in higher SNP errors, consistent with results reported by Zimin and Salzberg [55].

The rates of indel errors in the assemblies generated showed substantial differences across sequencing type, coverage, and polishing (**Figure 2G H**). Raw assemblies of Nanopore data contained the highest amount of indel errors with Illumina assemblies containing the least, both by significant margins. Raw hybrid assemblies contained low amounts of indel errors compared to the long-read assemblies, with raw PacBio assemblies having rates between Nanopore and hybrid. Coverage had a substantial effect on initial indel rates across Nanopore, PacBio, and hybrid assemblies, though this was particularly pronounced for Nanopore and PacBio data. With sufficient coverage, indel rates in PacBio assemblies were comparable to hybrid assemblies; however, the same was not true of Nanopore assemblies, which appeared to level out at a higher rate of indels. However, when polished, indel rates were substantially decreased across long-read and hybrid assemblies, indicating the benefit of incorporating Illumina data into assemblies. Indel rates in final Nanopore assemblies were slightly higher than PacBio or hybrid assemblies, but were at much more comparable levels. The amount of gain at this stage was affected by the polishing algorithm used, with NextPolish being particularly consistent across coverage levels at reducing indel errors (**Supplementary Figure S9**).

#### Computational performance

Illumina assembly methods, largely reliant on DBG-based approaches, tended to have the shorted wall-clock time in general. However, the fastest long-read and hybrid methods did out-speed them in some cases (**Supplementary Figure S10**). Long-read and hybrid methods showed a large range of wall-clock times. Of particular note is Canu, which took the longest of all methods considered by a substantial margin, especially when run on Nanopore data, likely due to its read correction step. This was still the case, even when Canu was run in -fast assembly in results graphs and outputs, indicating that Canu is not suited to a single-node computational architecture. Generally, time taken scaled with increasing coverage, an effect which was more pronounced for some assemblers than others. MaSuRCA-based methods in particular scaled well with increased coverage for both Illumina and hybrid data, though they tended to take longer initially than other assemblers. Assemblers such as w2rap, ABySS, Shasta, Raven, Miniasm, WTDBG2, and HASLR also tended to scale well with coverage.

All assemblers and polishing packages successfully completed using less than 128 GB of peak RAM, with most packages using less than 64 GB even at the highest coverage level (**Supplementary Figure S11**). Assemblers tended to increase in total memory usage with coverage, though this was less pronounced for some assemblers, such as HASLR and NECAT. A notable outlier was FALCON, which used the most memory for each coverage level; however, it appeared to peak close to the maximum available for 40X of coverage, and ran successfully at full coverage, indicating that this may be an optimising measure rather than a strict ceiling.

#### Ranking

Cumulative z-scores were calculated using N50 values, BUSCO completion percentages, BUSCOMP identity percentages, SNP error rates, and indel error rates for each assembly, comparing within each coverage level. The best polished assembly according to cumulative z-score, using any polisher or technology, for each assembler was extracted, and the best three of these assemblies for each data type are displayed in **Figure 4**, with the details for these assemblies provided in **Table 5**. For Illumina assemblies, SPAdes and MaSuRCA performed consistently well across coverage levels, though the overall quality of these assemblies was eclipsed by long-read and hybrid assemblies for higher coverage. Flye was a consistent top performer for Nanopore input data, creating the most contiguous assemblies with very minimal trade-off on BUSCO complete percentage and accuracy. Raven is also notable as a top performer, performing consistently across coverage levels. For PacBio data, Flye again results in the highest cumulative scores, in part due to its high contiguity of assembly. Canu performs well for higher coverage amounts, but takes considerably longer to run. For hybrid data, DBG2OLC performs extremely well for low coverage input, but scales poorly with increased coverage, and in some cases produces lower quality assemblies, indicating that high coverage levels may benefit from being subsampled before using this package. MaSuRCA and MaSuRCA+Flye are consistent in their overall cumulative z-scores across coverage levels, while HASLR, specifically run using PacBio data, performs well when given sufficiently high coverage data.

**Figure 4:**
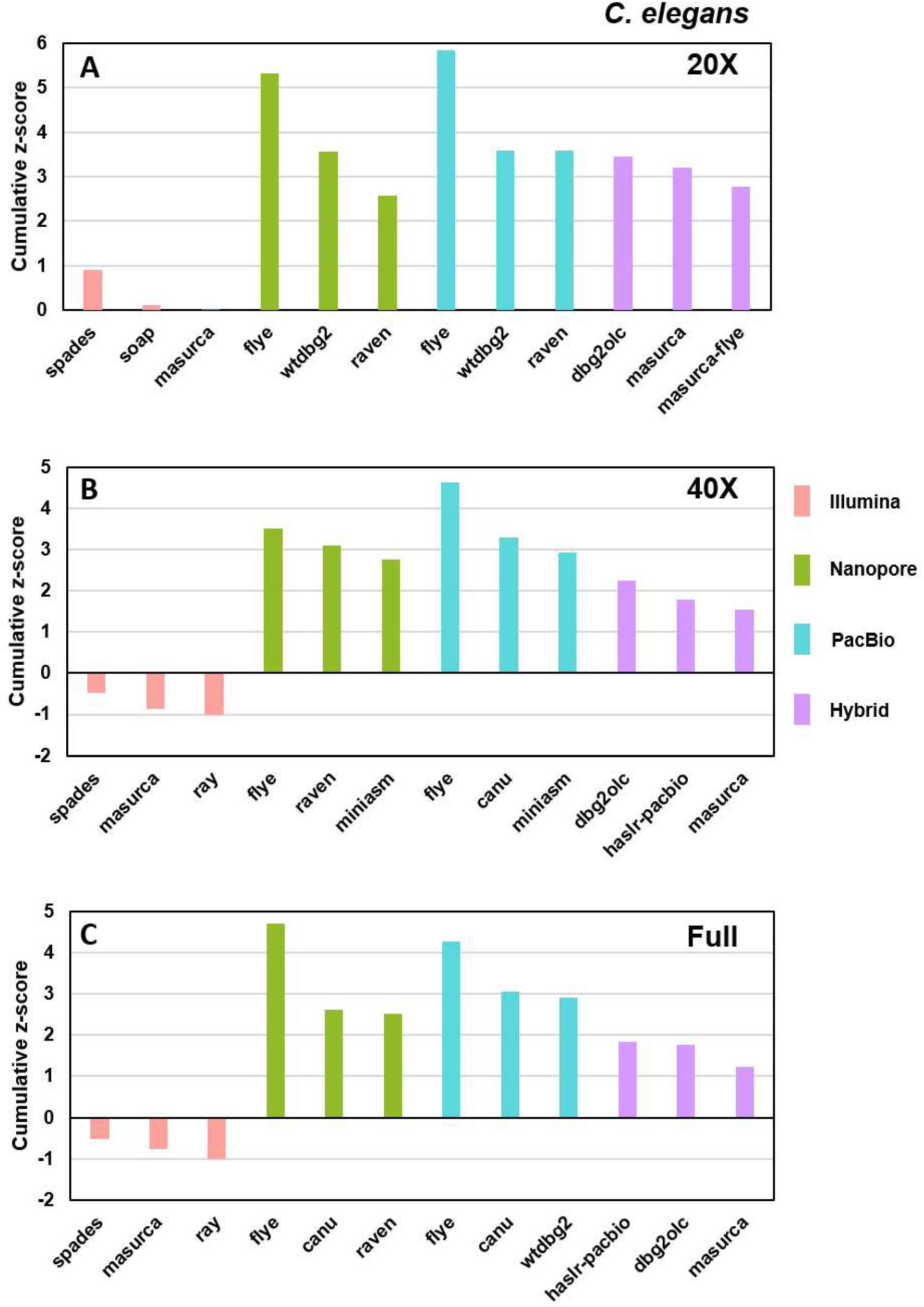
Plots of the cumulative z-scores for the three highest-scoring assemblies for each sequencing type, for each level of coverage, for *C. elegans*. Higher cumulative z-scores indicate better performance across multiple metrics, specifically N50, complete BUSCO percent, identical BUSCOMP percent, SNPs per 100 kbp, and indels per 100 kbp.

**Table 5:**
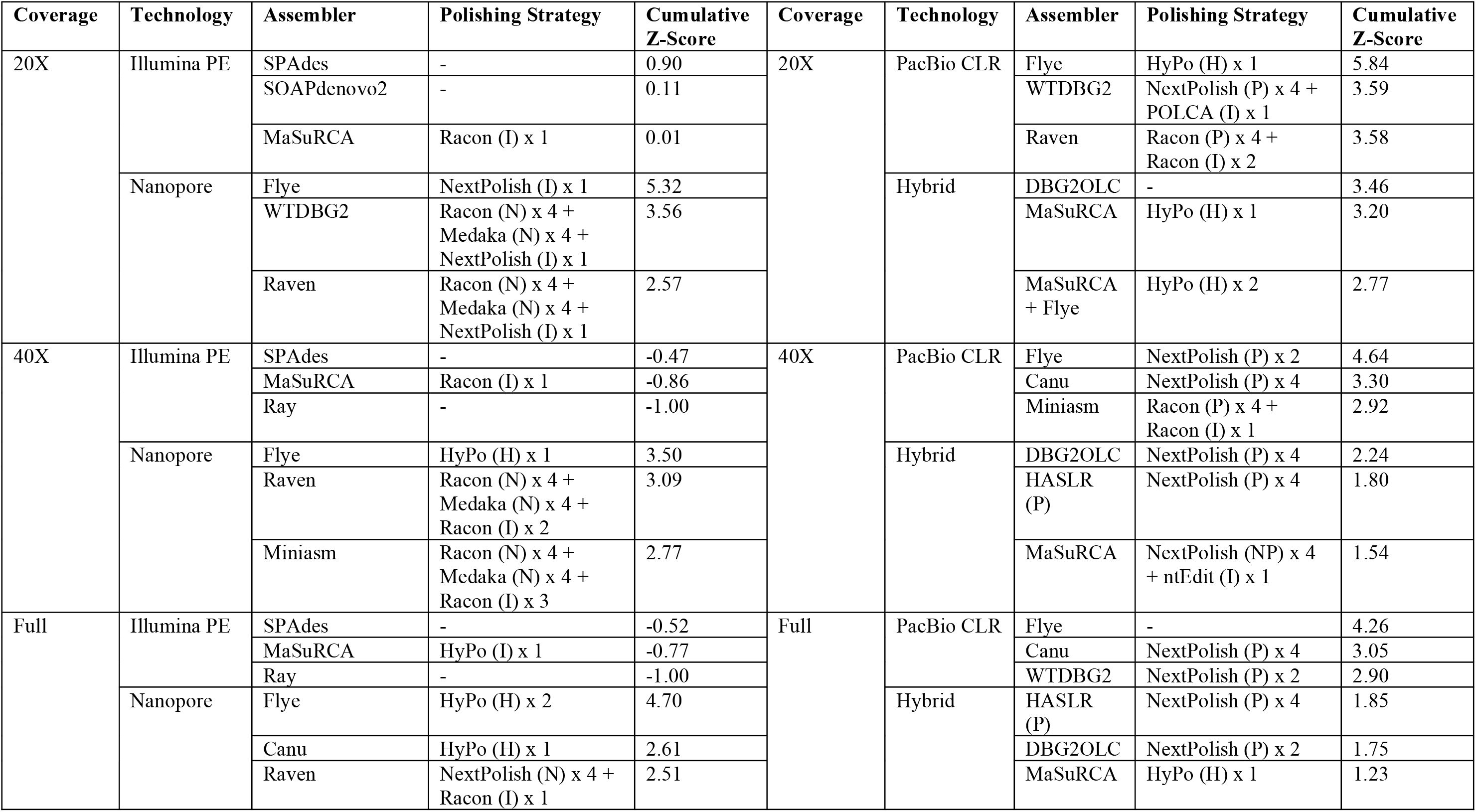
Details of best assemblies for *C. elegans* from each initial assembler, including details of polishing. (I) indicates Illumina data input, (N) indicates Nanopore data input, (P) indicates PacBio data input, and (H) indicates hybrid (short- and long-read) data.

### The D. melanogaster data set

*D. melanogaster* is a fruit fly which has been extensively used as a model organism across multiple disciplines [69 72]. Its genome was the second whole-genome sequenced for a multicellular organism [73], and has also been characterised to a substantial degree, with a plethora of genomic resources available [19]. The diploid reference genome assembly used in this study, *DmelR6* [73], is approximately 144 Mbp in size, distributed across three autosomes, two heterochromosomes, and a mitochondrial genome, and has a GC content of 42.1%, with a repeat content of 23.2% as measured by Red [59].

#### Contiguity and structural statistics

Contiguity did not change substantially between raw assemblies and polished ones, indicating as was the case for *C. elegans* that the contiguity of the assembly is dictated primarily by the assembly package (**Figure 5**). Again, long-read assembly methods produced the highest contiguity assemblies, with PacBio assemblies generally producing higher N50 values than Nanopore, with notable exceptions being assemblies generated using Nanopore data by Flye (**Figure 5C D**). Generally, N50 values increased with higher levels of coverage, with a particularly notable effect for PacBio assemblies. Illumina assemblies had substantially lower N50 values. Hybrid assemblies were generally comparable with long-read assemblies in N50 for low coverage, but fell behind for higher levels of coverage.

**Figure 5:**
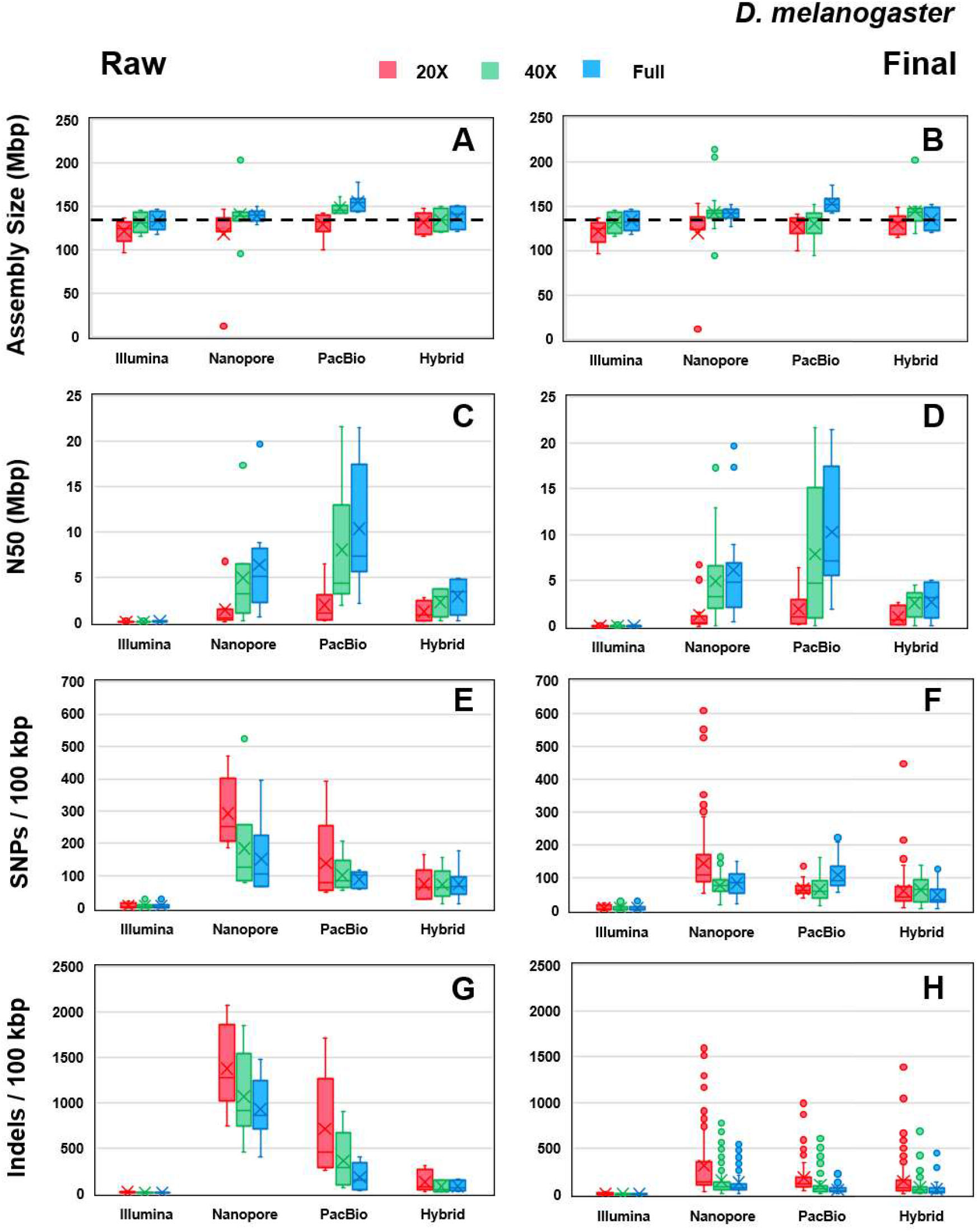
Box-and-whisker comparative plots for D. melanogaster for raw assemblies, i.e. without polishing, and final assemblies, i.e. polished with four rounds of long reads, followed by three rounds of short reads. lllumina assemblies are unpolished in both ‘raw’ and ‘final’ and are present for comparative purposes. The median is indicated by a horizontal line in each box, and the mean is indicated by a cross. Boxes represent the interquartile range of input assemblies. **Row 1:** Total assembly size for **(A)** raw and **(B)**final assemblies. **Row 2:** N50 for **(C)** raw and **(D)** final assemblies. **Row 3:** Average mismatch errors (SNPs) per 100 kbp of **(E)** raw and **(F)** final assembly sequence. **Row 4:** Average insertions and deletions per 100 kbp of **(G)** raw and **(H)** final assembly sequence.

The size of the assembly generated also did not increase with polishing, indicating that initial assembly primarily determines overall coverage of the genome (**Figure 5A B**). Assembly size generally increased with coverage, particularly in the case of PacBio assemblies. Long-read assemblies at higher coverage tended to overestimate the assembly size, as with *C. elegans* above. Hybrid assemblies were again very close to the reference size.

The GC% was underestimated slightly by long-read assemblers at the raw assembly stage, with Illumina assemblers tending to overestimate slightly, and hybrid assemblies producing assemblies which were in broad agreement with the reference (**Supplementary Figure S12**). After polishing, the final assemblies for Nanopore data moved toward the reference value more substantially than the PacBio assemblies.

At the raw assembly stage, assemblers across all input data types tended to overestimate the repeat content of the genome, particularly for Illumina assemblies, though Nanopore and PacBio assemblers showed a broad range of estimates (**Supplementary Figure S12**).

Polishing had a more substantial effect on total repeat length than for *C. elegans*, with Nanopore assemblies producing extremely close estimates in line with reference values after polishing, with notable improvement in PacBio estimates as well. Hybrid assemblies showed little change in repeat length with polishing. There was also minimal improvement in total repeat length as coverage increased.

#### Gene completeness

Illumina and hybrid assemblies for *D. melanogaster* produced high gene completeness with no polishing as measured by BUSCO, in many cases even with coverage levels as low as 20X (**Supplementary Figures S13 14**). Nanopore assemblies produced the lowest completion percentages at the raw assembly stage, with PacBio assemblies falling in between the two extremes. Generally, BUSCO complete percentages increased with coverage, with notable exceptions being the hybrid assembly packages. Some packages also show minimal improvement above 40X of coverage, indicating that, for the purposes of gene completeness, there may be an optimal level of coverage to achieve maximised BUSCO scores. Upon polishing, long-read assemblies made significant gains on the Illumina and hybrid scores, particularly at high coverage, with almost every assembler producing final assemblies with BUSCO completion percentages above 95% with at least one possible polishing combination. More variation is seen at low coverage levels, particularly for long-read assemblers, with the majority of the scores being significantly lower. However, there are assembly packages and polishing combinations able to produce assemblies with high gene completion, even at 20X of coverage, notably Flye, Raven and WTDBG2 for Nanopore data, and Flye for PacBio data. MaSuRCA, MaSuRCA+Flye and DBG2OLC assemblies also produced highly complete assemblies at 20X of coverage using hybrid data (**Supplementary Figure S15**), presenting multiple options for low-coverage sequencing projects where high gene completion is the goal.

Results from BUSCOMP analysis again indicate substantially higher gene completeness for the raw assemblies than BUSCO alone. In many cases, the BUSCOMP complete percentages are considerably above 90%, even at low coverage, with long-read assemblies being estimated substantially higher in completeness by BUSCOMP than BUSCO (**Figure 6C**). In addition, as for *C. elegans* above, BUSCO appears to give a good indication of the current state of the assembly, increasing with polishing up to a peak, while BUSCOMP appears to give a good indication of potential polished assembly quality, even at the raw assembly stage (**Figure 6D E**).

**Figure 6:**
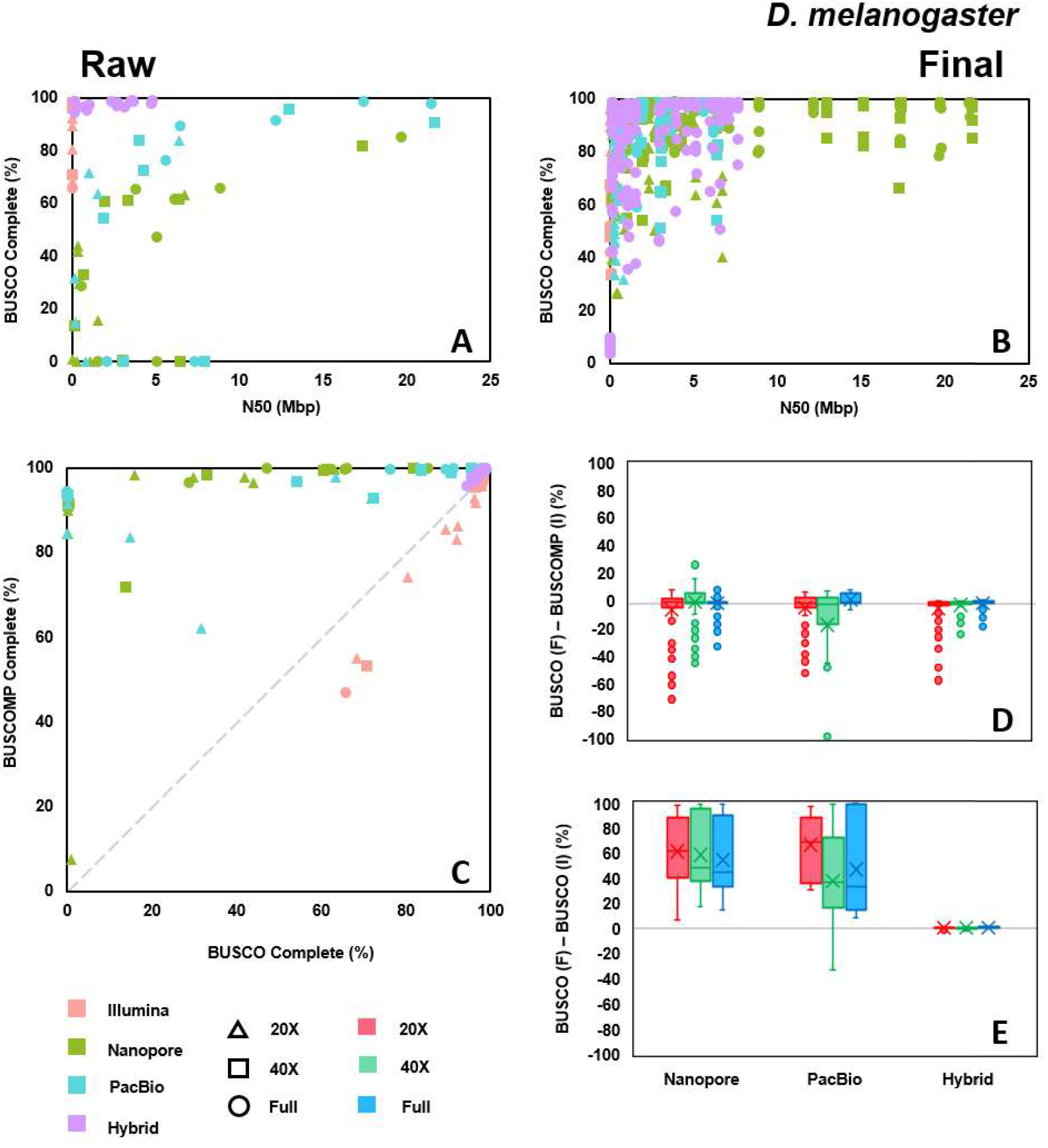
Comparison of complete BUSCO genes for D. melanogaster for raw assemblies, i.e. without polishing, and final assemblies, i.e. polished with four rounds of long reads, followed by three rounds of short reads. Illumina assemblies are unpolished in both ‘raw’ and ‘final’ and are present for comparative purposes. For the box-and-whisker plots, the median is indicated by a horizontal line in each box, and the mean is indicated by a cross. Boxes represent the interquartile range of input assemblies. **Top:** Percentage of complete BUSCO genes found in **(A)** raw and **(B)** final assemblies. **Bottom Left: (C)** Comparison of the percentage of complete BUSCOMP genes found in the raw assemblies against the percentage of BUSCO genes; the diagonal indicates equality between the two metrics. **Bottom Right:** Box-and-whisker plot comparisons by sequencing type of the difference between the percentage of **(D)** complete BUSCOMP genes or **(E)** complete BUSCO genes in the raw assembly, and the BUSCO genes in the final assembly, with the horizontal grey line indicating no change between raw and final. Negative values indicate overestimation by the raw assembly metric of the final assembly metric, while positive values indicate underestimation.

#### Accuracy

In a similar way to *C. elegans* above, the number of misassemblies did not prove to be a useful metric when comparing between assemblers, with the rates of SNP and indel errors being more robust to poor overall assembly coverage. At the raw assembly stage, similar results are seen for SNP errors for *D. melanogaster* as for *C. elegans*, with Nanopore assemblies having substantially higher SNP error rates, though slightly less pronounced, and Illumina assemblies having the lowest (**Figure 5E F**). SNP errors tend to decrease with coverage across long-read and hybrid assemblies at the raw assembly stage. Polishing tends to improve Nanopore, PacBio and hybrid assemblies with respect to SNP errors, with the effect most pronounced for Nanopore and PacBio data, particularly at low coverage levels. As with *C. elegans*, there is a minor increase in SNP error rates with polishing for high-coverage PacBio data. Short-read polishing with HyPo, Pilon, and POLCA performed consistently well across all data types, with small additional improvements compared to NextPolish and Racon in this case (**Supplementary Figure S18**).

The indel error rates for the raw assemblies showed similar trends in *D. melanogaster* as in *C. elegans*, with long-read- and hybrid-based assemblies having considerably higher indel error rates compared to Illumina (**Figure 5G H**). Again, Nanopore assemblies had the highest rates of indel errors, followed by PacBio. Increasing coverage did tend to improve indel error rates across long-read and hybrid assemblies, though this was less pronounced for Nanopore assemblies compared to PacBio assemblies, reaching a higher ‘minimum’ rate. Polishing however does considerable work for long-read and hybrid assemblies, with indel rates reduced considerably, particularly for Nanopore assemblies. Short-read polishing with NextPolish tended to have the most consistent impact on reducing indel errors across coverage levels, though a range of short-read polishers also performed comparably well, particularly at higher coverage levels (**Supplementary Figure S20**)

#### Computational performance

The speed of assembly across short- and long-read assemblers was closer for *D. melanogaster* than for *C. elegans*. While short-read assemblers did tend to produce fast assemblies for low coverage, these tended to scale more poorly for high coverage, with the majority of Nanopore and PacBio assemblers being comparable in speeds for high coverage (**Supplementary Figure S21**). Of note are the high-speed assemblers for long-read and hybrid data, namely Shasta, Raven, Miniasm, WTDBG2, and HASLR, which were all comparable to, or faster than, the fastest Illumina assemblers across coverage levels. Across the board, time generally scaled with increasing coverage, though as for *C. elegans*, this was more pronounced for some assemblers than others. Assemblers which scaled well with coverage were notably w2rap, SOAPdenovo2, and ABySS for Illumina data, Shasta and WTDBG2 for Nanopore data, WTDBG2 for PacBio data, and HASLR for hybrid data. Three assemblers were notable for extremely long run times, namely Canu, Falcon, and DBG2OLC. As before, Canu is not suited to a single-node computing environment, taking multiple days to run for both long-read data types, but particularly long for Nanopore, exacerbated by high coverage. Running Canu in ‘fast’ mode, indicated by canu-fast in results and plots, did improve this issue to some degree, but was still considerably slower than all other Nanopore assemblers. This poor coverage scaling was also evident for DBG2OLC, where, for 40X coverage, the assembly did not complete in the 5 day window allocated, indicating that smaller coverage, particularly of long reads, is likely more optimal for this particular assembler.

Peak memory usage varied across assemblers and sequencing types, but did not exceed the 128 GB allocation in any case, with the majority of assemblers completing with under 64 GB usage at peak (**Supplementary Figure S22**). For some assemblers, this memory usage was even below 8 GB specifically ABySS, Meraculous, SparseAssembler, Shasta, and HASLR making assembly on a middle-range laptop extremely possibility, albeit with increased wall clock time due to fewer than 20 cores. Platanus and FALCON were particularly noteworthy for requiring larger amounts of memory at peak, with FALCON again using almost the all allocated memory for the highest coverage levels, though as above, this may be more an optimisation measure than a ceiling, as this limit did not change considerably from 40X coverage to full, despite a jump of almost 60X of added coverage.

#### Ranking

Cumulative z-scores were calculated as for *C. elegans*, using N50 values, BUSCO completion percentages, BUSCOMP identity percentages, SNP error rates, and indel error rates for each assembly, compared within each coverage level. The best polished assembly, using any polisher or technology, for each assembler was extracted, and the best three of these assemblies for each data type are displayed in **Figure 7**, with additional details provided for each assembly in **Table 6**. For Illumina assemblies, SPAdes performed consistently well across all levels of coverage, with almost all other assemblers performing comparably well at higher coverage.

**Figure 7:**
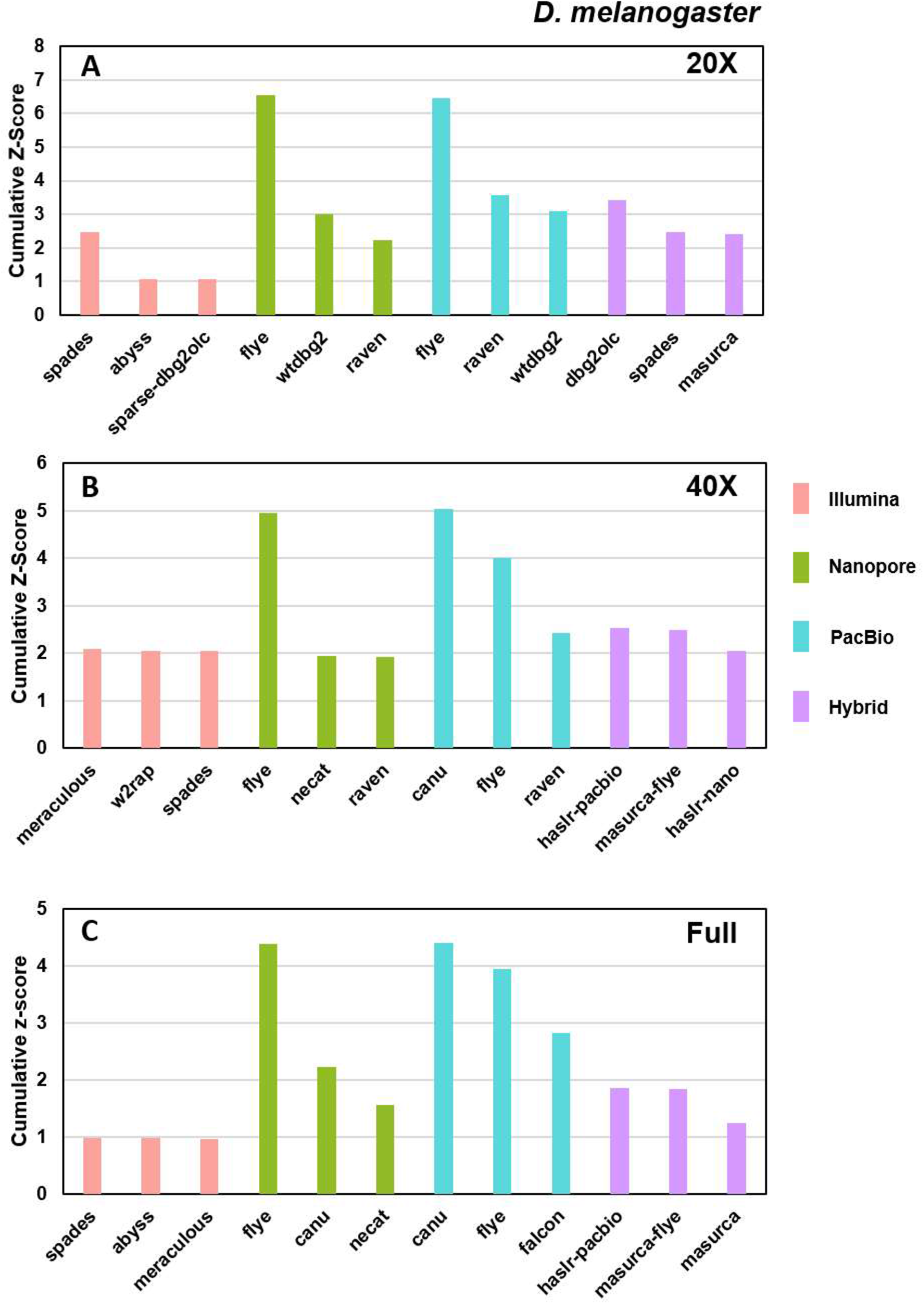
Plots of the cumulative z-scores for the three highest-scoring assemblies for each sequencing type, for each level of coverage, for *D. melanogaster*. Higher cumulative z-scores indicate better performance across multiple metrics, specifically N50, complete BUSCO percent, identical BUSCOMP percent, SNPs per 100 kbp, and indels per 100 kbp.

**Table 6:** Details of best assemblies for D. melanogaster from each initial assembler, including details of polishing. (I) indicates Illumina data input, (N) indicates Nanopore data input, (P) indicates PacBio data input, and (H) indicates hybrid (short- and long-read) data.

| Coverage | Technology | Assembler | Polishing Strategy | Cumulative Z-Score | Coverage | Technology | Assembler | Polishing Strategy | Cumulative Z-Score |
| --- | --- | --- | --- | --- | --- | --- | --- | --- | --- |
| 20X | Illumina PE | SPAdes | HyPo (I) x 2 | 2.47 | 20X | PacBio CLR | Flye | POLCA (I) x 3 | 6.45 |
|  |  | ABYSS | Racon (I) x 3 | 1.08 |  |  | Raven | NextPolish (P) x 4 + Pilon (I) x 3 | 3.58 |
|  |  | SparseAssembler + DBG2OLC | Racon (I) x 3 | 1.06 |  |  | WTDBG2 | NextPolish (P) x 4 + POLCA (I) x 3 | 3.09 |
|  | Nanopore | Flye | HyPo (H) x 4 | 6.56 |  | Hybrid | DBG2OLC (IP) | POLCA (I) x 3 | 3.43 |
|  |  | WTDBG2 | NextPolish (N) x 4 + NextPolish (I) x 1 | 3.02 |  |  | SPAdes (INP) | NextPolish (NP) x 4 + POLCA (I) x 3 | 2.47 |
|  |  | Raven | NextPolish (N) x 4 + NextPolish (I) x 1 | 2.22 |  |  | MaSuRCA (INP) | ntEdit (I) x 2 | 2.41 |
| 40X | Illumina PE | Meraculous | POLCA (I) x 3 | 2.10 | 40X | PacBio CLR | Canu | POLCA (I) x 1 | 5.03 |
|  |  | w2rap | POLCA (I) x 1 | 2.05 |  |  | Flye | HyPo (H) x 1 | 4.01 |
|  |  | SPAdes | GATK (I) x 2 | 2.05 |  |  | Raven | Racon (P) x 4 + HyPo (I) x 1 | 2.43 |
|  | Nanopore | Flye | HyPo (H) x 2 | 4.97 |  | Hybrid | HASLR (P) | HyPo (H) x 1 | 2.54 |
|  |  | NECAT | HyPo (H) x 2 | 1.94 |  |  | MaSuRCA + Flye (INP) | HyPo (H) x 4 | 2.48 |
|  |  | Raven | NextPolish (N) x 4 + Pilon (I) x 3 | 1.91 |  |  | HASLR (N) | HyPo (H) x 1 | 2.04 |
| Full | Illumina PE | SPAdes | GATK (I) x 2 | 0.98 | Full | PacBio CLR | Canu | Pilon (I) x 1 | 4.41 |
|  |  | ABYSS | Pilon (I) x 3 | 0.97 |  |  | Flye | Flye (I) x 1 | 3.95 |
|  |  | Meraculous | POLCA (I) x 1 | 0.96 |  |  | FALCON | HyPo (H) x 1 | 2.82 |
|  | Nanopore | Flye | HyPo (H) x 3 | 4.38 |  | Hybrid | HASLR (P) | HyPo (H) x 1 | 1.85 |
|  |  | Canu | HyPo (H) x 1 | 2.22 |  |  | MaSuRCA + Flye (INP) | HyPo (H) x 3 | 1.84 |
|  |  | NECAT | HyPo (H) x 3 | 1.56 |  |  | MaSuRCA (INP) | HyPo (H) x 4 | 1.24 |

As for *C. elegans* above, Flye produced consistently impressive assemblies using Nanopore data, with extremely high contiguity, even at low coverage levels while Raven was also notable for consistency across even low coverage. For PacBio data, Flye was again a consistently high performer, though the assemblies produced by Canu at high coverage levels did overtake these. DBG2OLC performed extremely well for low coverage of data, but fell off for higher levels compared to other hybrid assemblers. MaSuRCA+Flye and HASLR, specifically run on PacBio data, were reliable performers at high coverage, but did not overtake the highest performing long-read-only assembly methods. HASLR is particularly noteworthy in this context, as it had the shortest assembly time of any assembler tested, yet produced the best overall results among hybrid assemblers given sufficient coverage.

## Discussion

The options available to researchers for assembling novel genomes *de novo* are staggering. In order to help provide some guidance and data for informed decision making, we have presented key evaluation metrics for 14,000 genome assemblies, totalling 30 different raw assembly options, four long-read polishing options, seven short-read polishing options, and one hybrid polishing option. By comprehensively comparing Illumina, Oxford Nanopore, and PacBio CLR sequencing methods, as well as hybrid methods incorporating multiple of these, clear trends emerge, with no one assembler, polisher, or technology providing maximal values in every category. While results are largely consistent across the two genomes considered here, it is first and foremost important that multiple assemblers be trialled for novel organisms, in order to provide within-species comparisons as assembly progresses, to achieve the best possible outcome. This is further underscored by the close grouping of methods ‘at the top’, with multiple combinations of technologies, assemblers, and polishers all producing high quality results overall. Using even a small subset of the consistently high performers demonstrated here is likely to produce high quality assemblies in most cases.

With regard to sequencing technologies more specifically, there are some clear trends for both *C. elegans* and *D. melanogaster*. As has also been previously indicated by recent long-read sequencing efforts [74 75], long-read and hybrid assemblies produce consistently more contiguous assemblies than using only short reads. While N50 alone is not likely to dictate absolute assembly quality or gene content, the resolution of larger-scale structural variants and repeat regions is likely to be made easier for longer fragments of assembly [76]. In addition, when scaffolded using additional methods such as Hi-C, these assemblies may end up producing less gaps or misassemblies [64].

Both Nanopore and PacBio data assembled into highly contiguous assemblies. While raw Nanopore assemblies were generally of lower gene completeness and accuracy than PacBio raw assemblies, this difference is almost completely removed by polishing. When final assemblies are considered, Nanopore and PacBio both produce assemblies of comparable contiguity, gene completeness, and accuracy, with most tools taking similar time and memory consumption on either sequencing type.

Hybrid assemblies produced the most consistent raw assemblies; however, they were also consistently overtaken in final metrics by long-read assemblies after polishing. This suggests that producing an assembly from longer reads, even with lower accuracy, sets a firm foundation for final assembly quality, and that shorter, more high quality reads can be most useful to improve this scaffold once created.

There are also clear trends across assembly algorithms, with particular packages being consistent high performers, and other packages requiring specific conditions to perform best. For Illumina assemblies, SPAdes was a consistent top performer across both *C. elegans* and *D. melanogaster* across multiple metrics, with notable other strong performers being MaSuRCA, for gene completeness, and w2rap, for speed.

For Nanopore assemblies, Flye was a standout performer, producing highly contiguous assemblies with extremely high completeness and accuracy. Raven also performed well, but benefits considerably from polishing. Shasta performed well in accuracy and gene completeness, but only when given high levels of coverage; at low coverage levels such as 20X, Shasta failed to construct large portions of the reference genome in both cases. Canu gave good results for Nanopore data when given high levels of coverage, but was computationally the most expensive by a wide margin. However, Canu is notably designed to maximise usage of a cluster environment, and would likely be more reasonable in run-times when given multiple nodes of a high-performance computing system to run on. For PacBio assemblies, Flye again produced exceptional results across major metrics, with Canu being a more competitive alternative for PacBio than for Nanopore, with almost all assemblers producing final assemblies of reasonable quality. FALCON is notable for producing high quality assemblies when given sufficient coverage; however, it scales more poorly than other assemblers, taking considerably longer to run than any other PacBio assembler, with the exception of Canu. Hybrid assemblies were mixed. Lightning-fast assembly with hybrid methods is possible using HASLR, particularly with PacBio data, and benefits from higher coverage input data. MaSuRCA and MaSuRCA+Flye were consistently high-quality performers, particularly on gene completeness and accuracy, but took considerably longer to run.

Polishing of assemblies, particularly those generated by long-read sequencing alone, is almost always essential to producing assemblies of the highest quality across all metrics. The best polishing methods identified for each assembler were a broad range, but in almost all cases included short-read polishing in some form, with the best combination of methods for each assembler across all coverage levels for *D. melanogaster* using short reads, and all but the highest coverage PacBio assemblies for *C. elegans* using short reads. Given the significantly lower price point for short-read sequencing, there is substantial benefit to supplementing any long-read project with short-read sequencing.

Some differences do emerge between *C. elegans* and *D. melanogaster*. First, the Illumina assemblies for *C. elegans* are of considerably lower quality than for *D. melanogaster*. Whether this is due to a peculiarity of the *C. elegans* genome or the particular set of *C. elegans* data used is unknown. In addition, the estimation of GC% and repeat length is inconsistent between both genomes, with long-read assemblers overestimating or underestimating depending on the organism.

Coverage is an important consideration before commencing any assembly project, and the data presented here will likely impact upon these decisions. By looking at 20X, 40X, and a ‘full’ set of coverage (>40X), there are clear benefits to high coverage sequencing. Assembly size is higher with higher coverage, implying more thorough assembly across the genome; N50 values also increase with coverage. Higher coverage assemblies also typically produced higher accuracy in the final assembly, with lower SNP and indel error rates. However, these differences were more pronounced for some assemblers than others. For long-read data, Flye was a consistent high-performer for low-coverage data, along with SPAdes for Illumina data and DBG2OLC for hybrid data, each generating highly complete assemblies with reasonably high accuracy. In addition for long-read assemblies, polishing with a hybrid strategy either using short- and long-reads together through HyPo, or by polishing first with long reads using Medaka for Nanopore, or NextPolish or Racon for PacBio, followed by short reads using NextPolish or POLCA is also successful, even at low coverage. While this many not hold for genomes with particularly complex regions of repeats, or other distinct features making assembly difficult, it is likely that many assembly projects can get by even with a small sequencing budget and still produce an assembly of high quality.

This work presents an extensive evaluation across almost all commonly used tools for genome assembly and polishing in recent literature. Given the scale of this undertaking, there are therefore considerable constraints and limitations that should be addressed. We have chosen here to benchmark on organisms meeting certain criteria, namely the existence of a gold-standard genome reference assembly constructed using input from a technology not considered here such as Sanger sequencing, that also has sufficient data publicly available to benchmark across coverage levels for all three technologies. In addition, given the sheer scale of combinations and computational time required to complete this benchmarking, we were also constrained in the total genome size to be considered, with assemblies of both mouse and human, both also meeting the above criteria, significantly more computationally intensive. As time progresses and more model genomes are inevitably sequenced and re-sequenced, it will likely be possible to expand this evaluation to additional organisms and data types. Limitations were also placed on the polishing of assemblies, with strict numbers of rounds selected (four rounds for long reads, three rounds for short reads), including when combining long-read and short-read polishing, with the four-times-polished assembly always chosen regardless of comparative quality, in order to manage the sheer scale of assemblies being generated. Further investigation into optimal polishing strategies is still required.

Given the trend toward chromosome-level assembly, it is also noted that we have not looked at the effect of scaffolding methods such as Hi-C or BioNano, in part due to sourcing of data, and in part due to sheer combinatorial size of methods already under consideration. Additional pre- and post-processing methods, such as data cleaning, error correction, haplotype reduction, and post-assembly screening, are also likely to have considerable impacts upon final assembly quality. We note that recent efforts have begun to benchmark these methods for a selection of tools and technology inputs [15 17]. We hope that our contribution to interrogating and evaluating methods across the broad spectrum available aids in spurring additional benchmarking work to target these further steps.

## Key Points

- Oxford Nanopore and PacBio CLR assemblies, when combined with short-read Illumina data for polishing, produce the highest quality assemblies across multiple metrics. Hybrid assemblies produce consistent results but have lower peak performances. Illumina-only assemblies are more accurate at a per-base level, but are more fragmented by multiple orders of magnitude.
- Peak performers, when weighting key metrics equally, were SPAdes and MaSuRCA for Illumina assemblies, Flye for Oxford Nanopore and PacBio data, and DBG2OLC and HASLR for hybrid assemblies.
- Polishing is essential for long-read assemblies, but less so for hybrid and Illumina assemblies. Hybrid polishing strategies with HyPo produce consistently high results for Oxford Nanopore and PacBio; long-read polishing followed by short-read polishing also produces high results; short-read-only polishing is sufficient for the best raw PacBio assemblies, such as those produced by Flye or Canu.
- Coverage of sequencing data can improve the likelihood of a high-quality assembly; however, 20X of coverage of long reads of either type with 20X of coverage of short reads may be sufficient to still produce comparable quality while saving considerably on both time and money, though this may change for more difficult to assemble genomes.

## Acknowledgments

The authors would like to thank the team at CSIRO Scientific Computing for their help in troubleshooting the many algorithmic packages used in this work.

## Data availability

The code for the benchmarking study is available at https://github.com/genomeassembler/benchmarking-study.

