## Supplementary Files for "Exhaustive benchmarking of *de novo* assembly methods for eukaryotic genomes"

### SUPPLEMENTARY MATERIAL

#### 1 Supplementary Figures

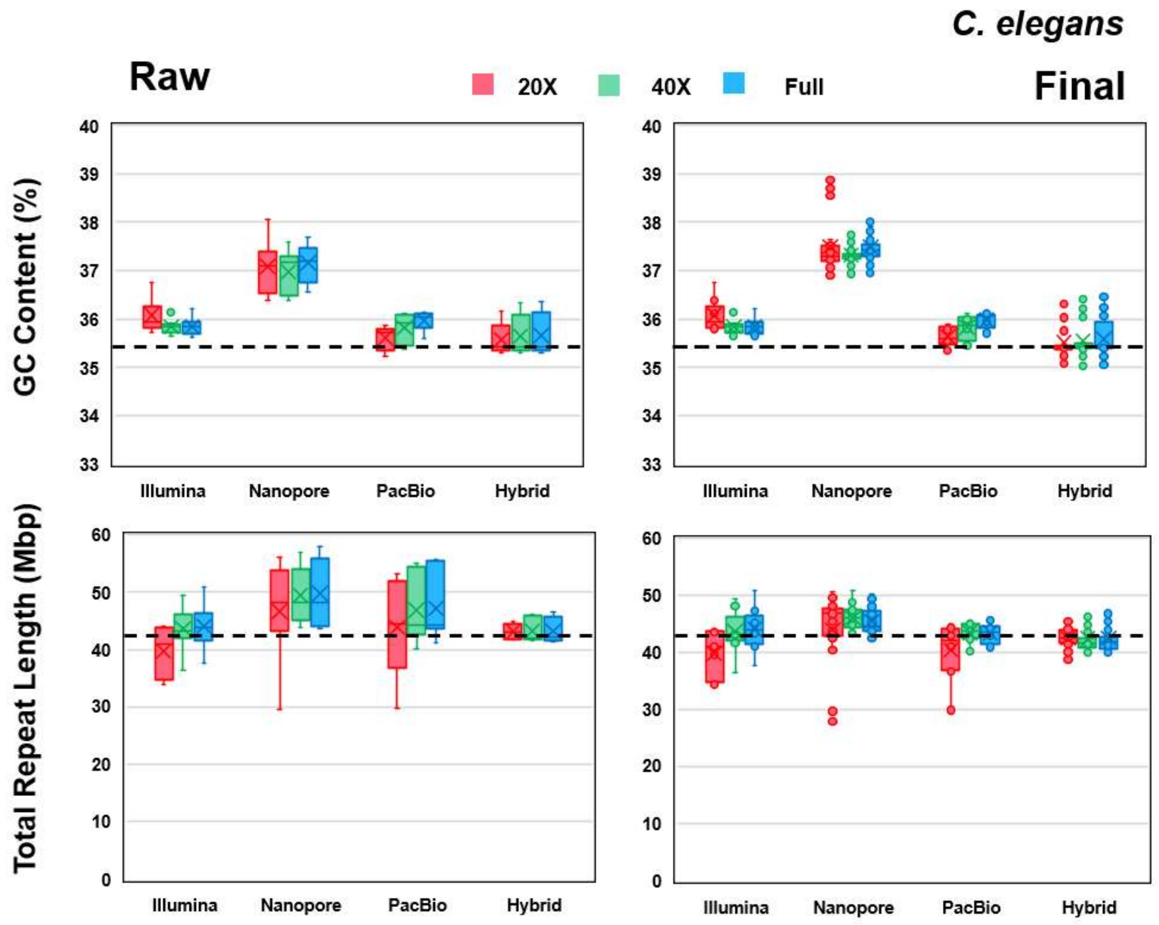


**Supplementary Figure S1:** Additional box-and-whisker plots of contiguity metrics for *C. elegans*, comparing GC% and total repeat length between raw and final assemblies. The dotted horizontal lines indicate the reference values.

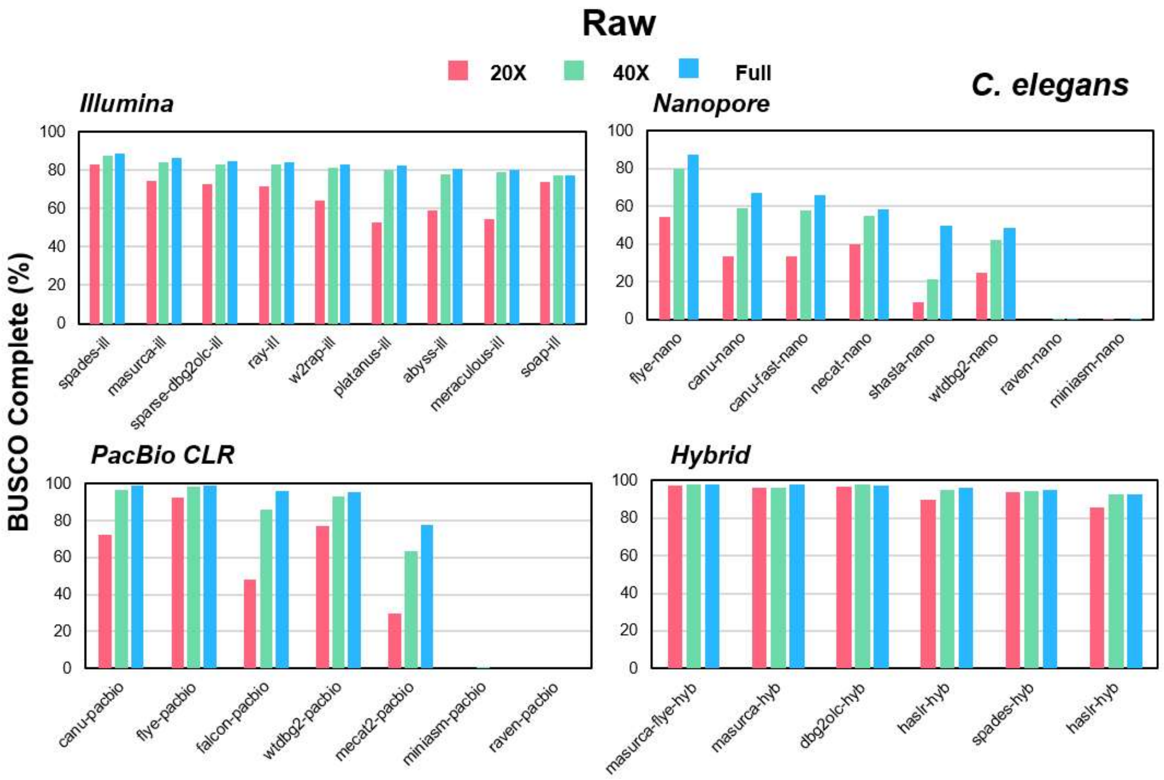


**Supplementary Figure S2:** BUSCO complete percentages for all raw assemblies for *C. elegans* at 20X (red), 40X (green) and full (blue) levels of coverage.

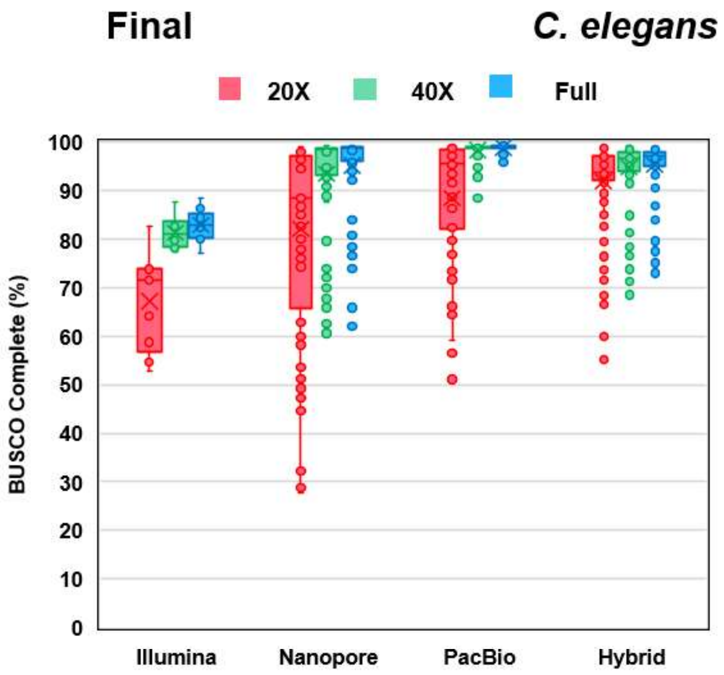


**Supplementary Figure S3:** Box-and-whisker plots of BUSCO complete percentages for all final assemblies for *C. elegans* at 20X (red), 40X (green) and full (blue) levels of coverage, by sequencing type.

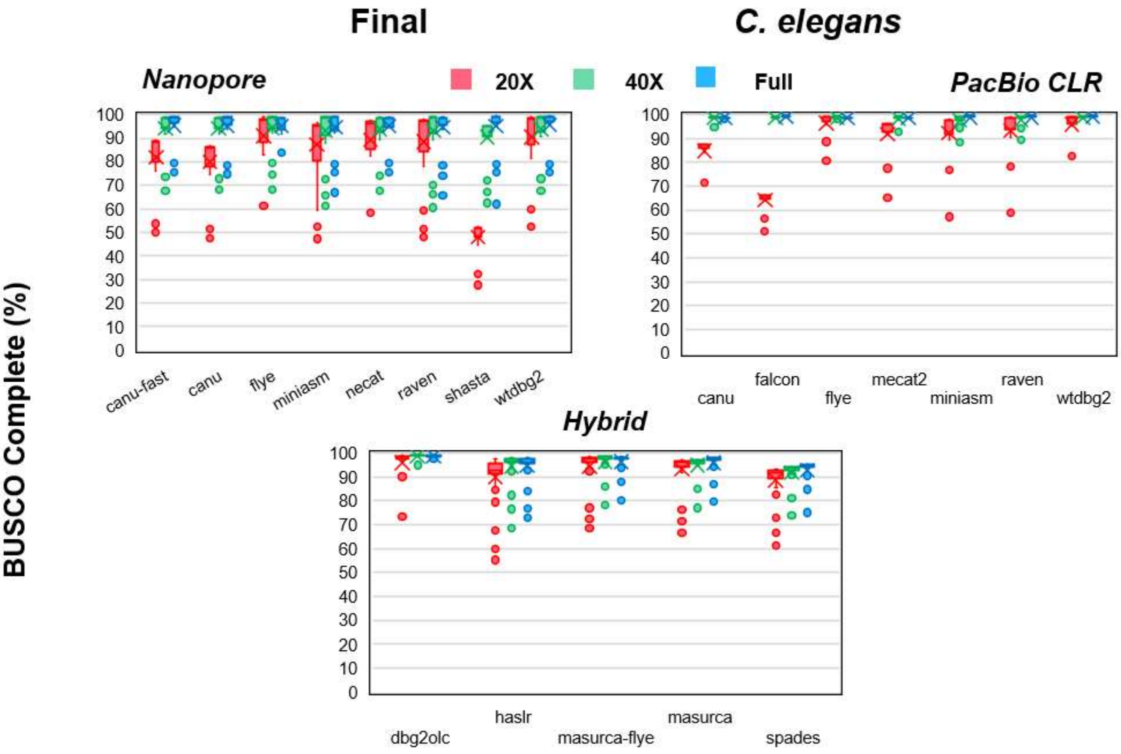


**Supplementary Figure S4:** Box-and-whisker plots of BUSCO complete percentages for all final assemblies for *C. elegans* at 20X (red), 40X (green) and full (blue) levels of coverage, by assembler.

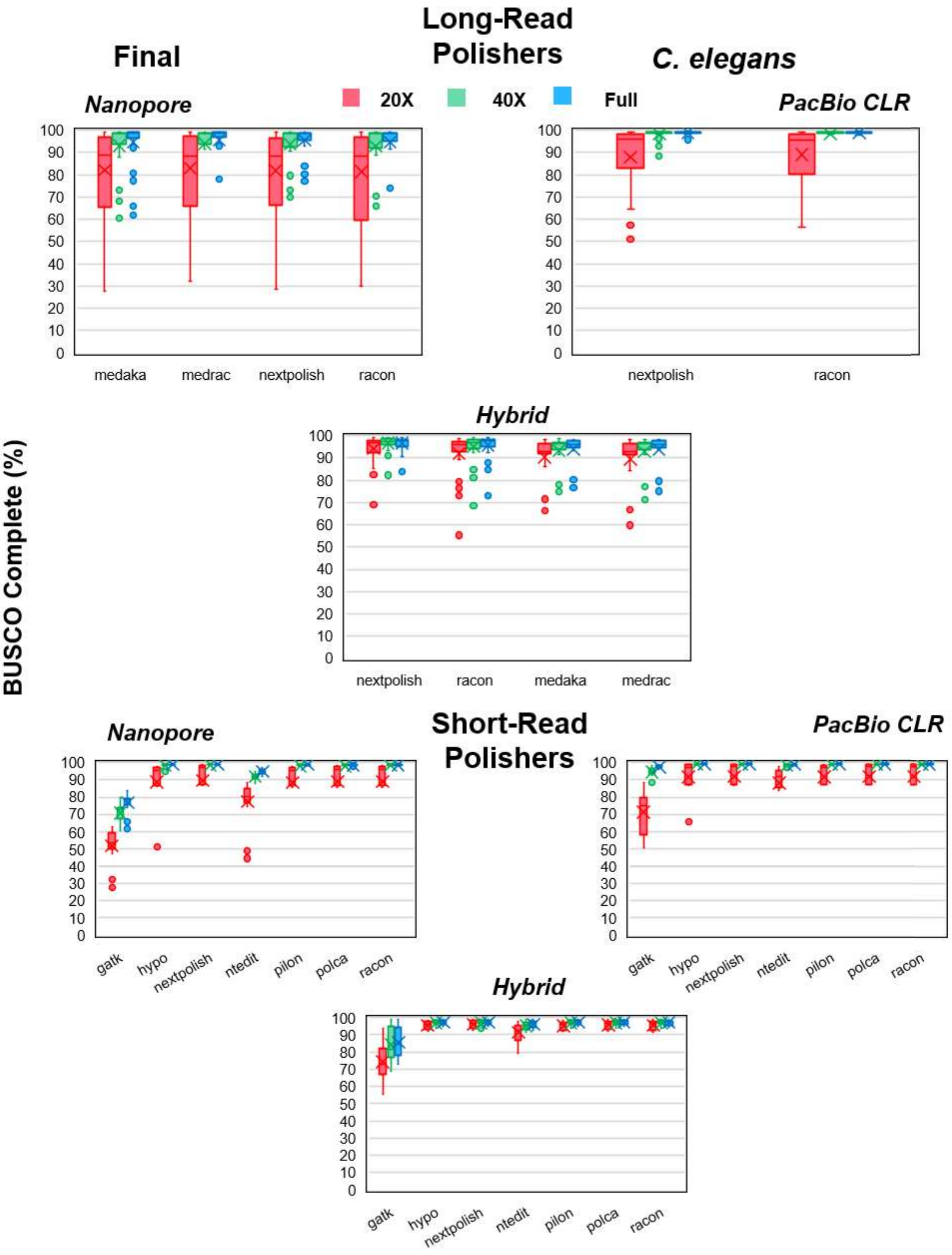


**Supplementary Figure S5:** Box-and-whisker plots of BUSCO complete percentages for all final assemblies for *C. elegans* at 20X (red), 40X (green) and full (blue) levels of coverage, by polishing algorithm used.

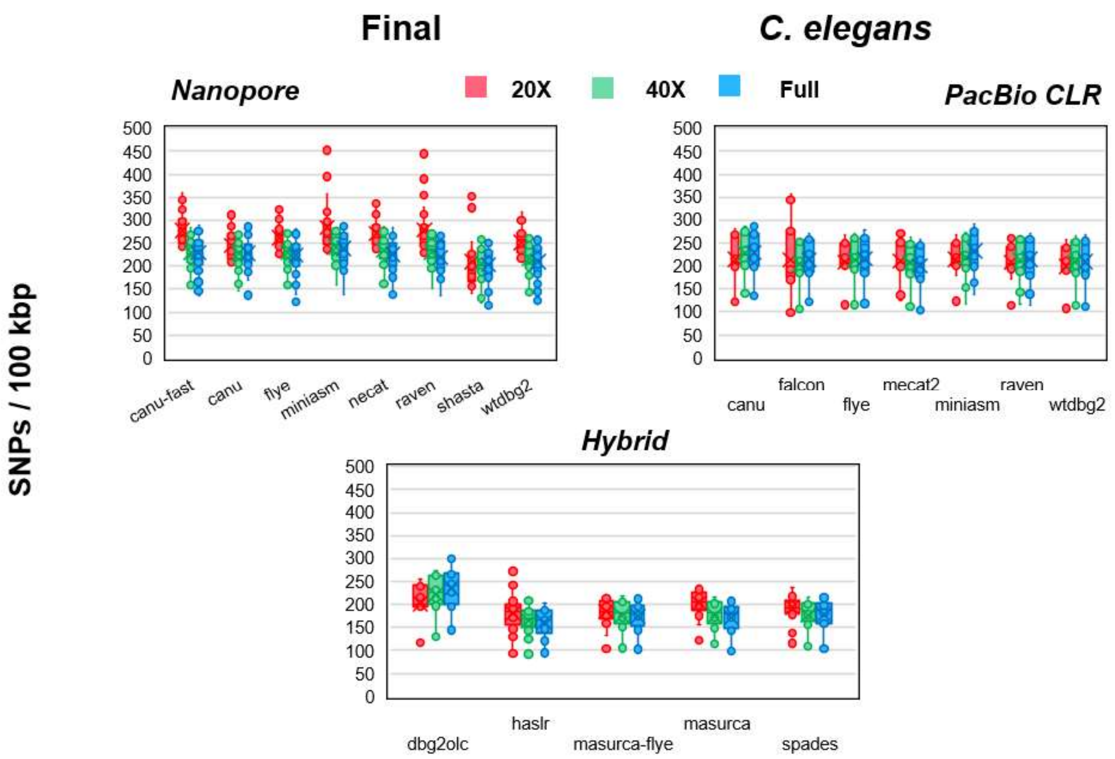


**Supplementary Figure S6:** Box-and-whisker plots of SNP error rates per 100 kbp of sequence for all final assemblies for *C. elegans* at 20X (red), 40X (green) and full (blue) levels of coverage, by assembler.

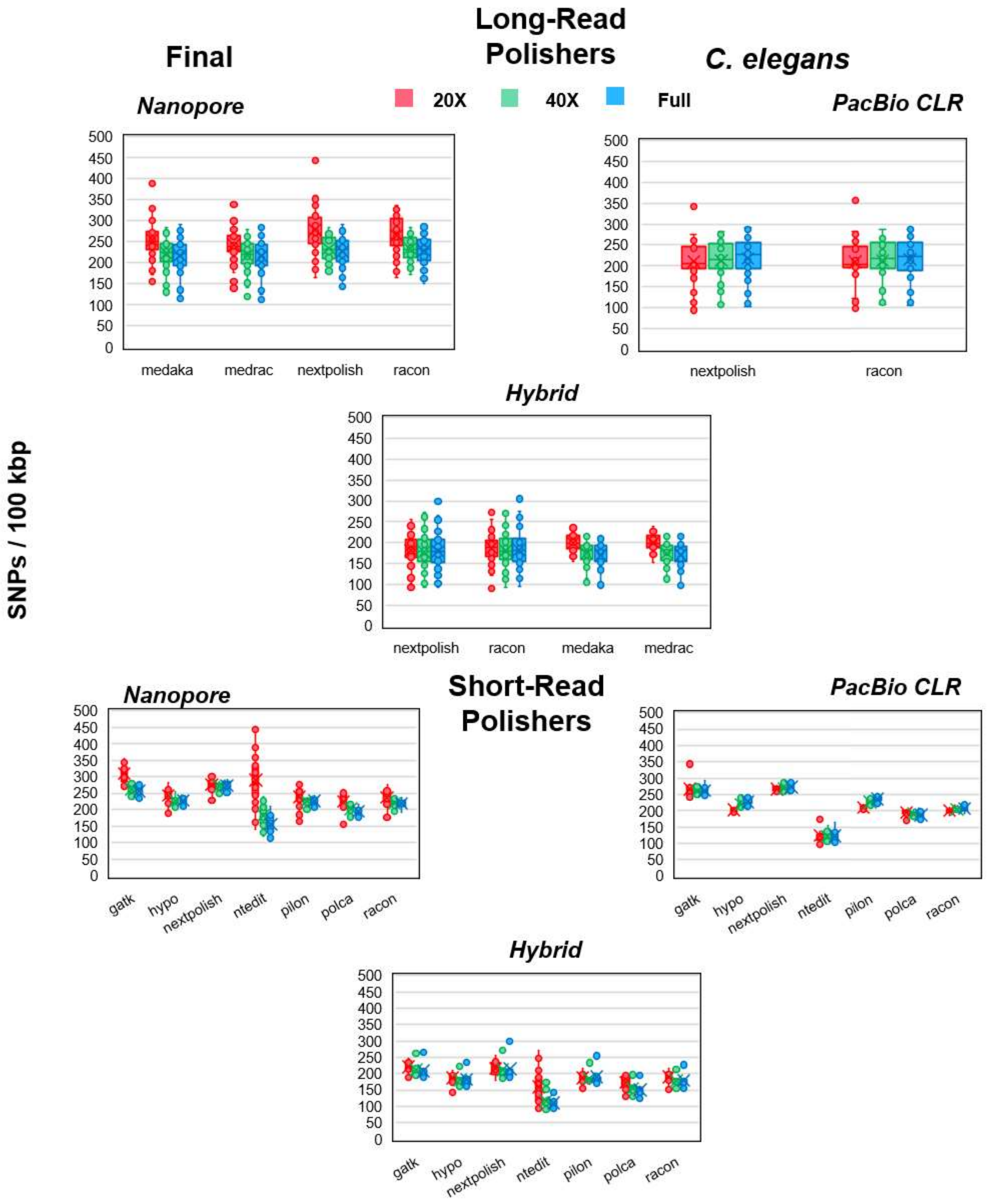


**Supplementary Figure S7:** Box-and-whisker plots of SNP error rates per 100 kbp of sequence for all final assemblies for *C. elegans* at 20X (red), 40X (green) and full (blue) levels of coverage, by polishing algorithm used.

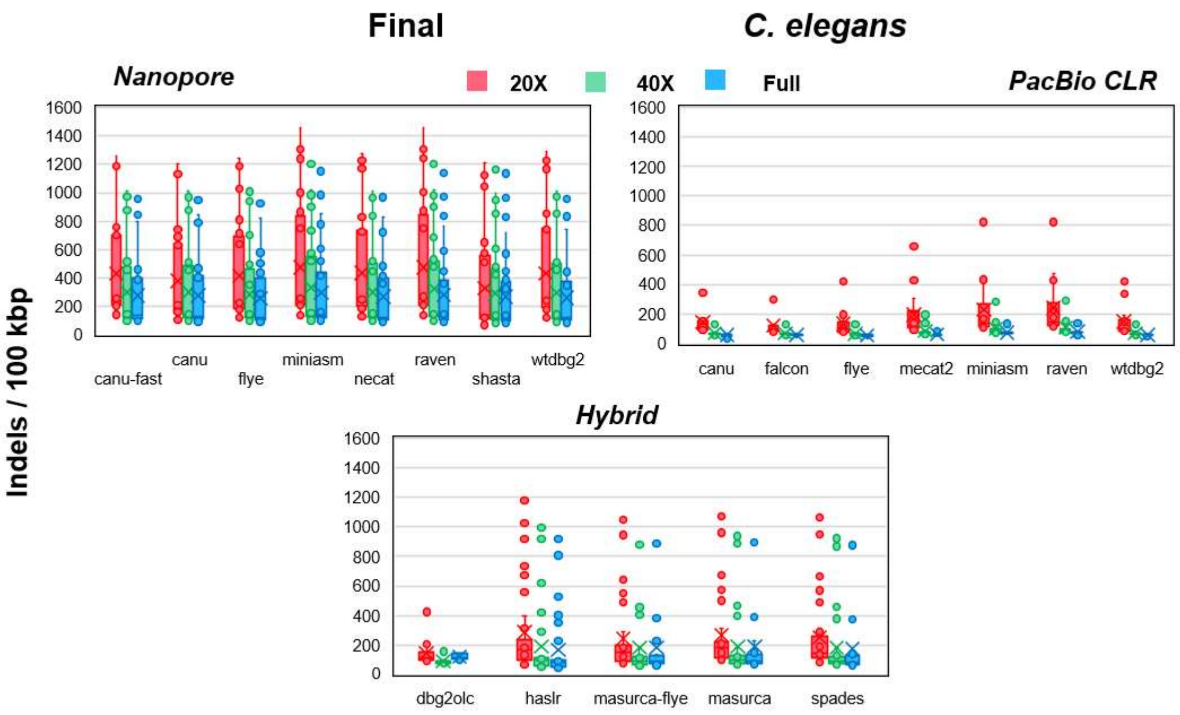


**Supplementary Figure S8:** Box-and-whisker plots of indel error rates per 100 kbp of sequence for all final assemblies for *C. elegans* at 20X (red), 40X (green) and full (blue) levels of coverage, by assembler.

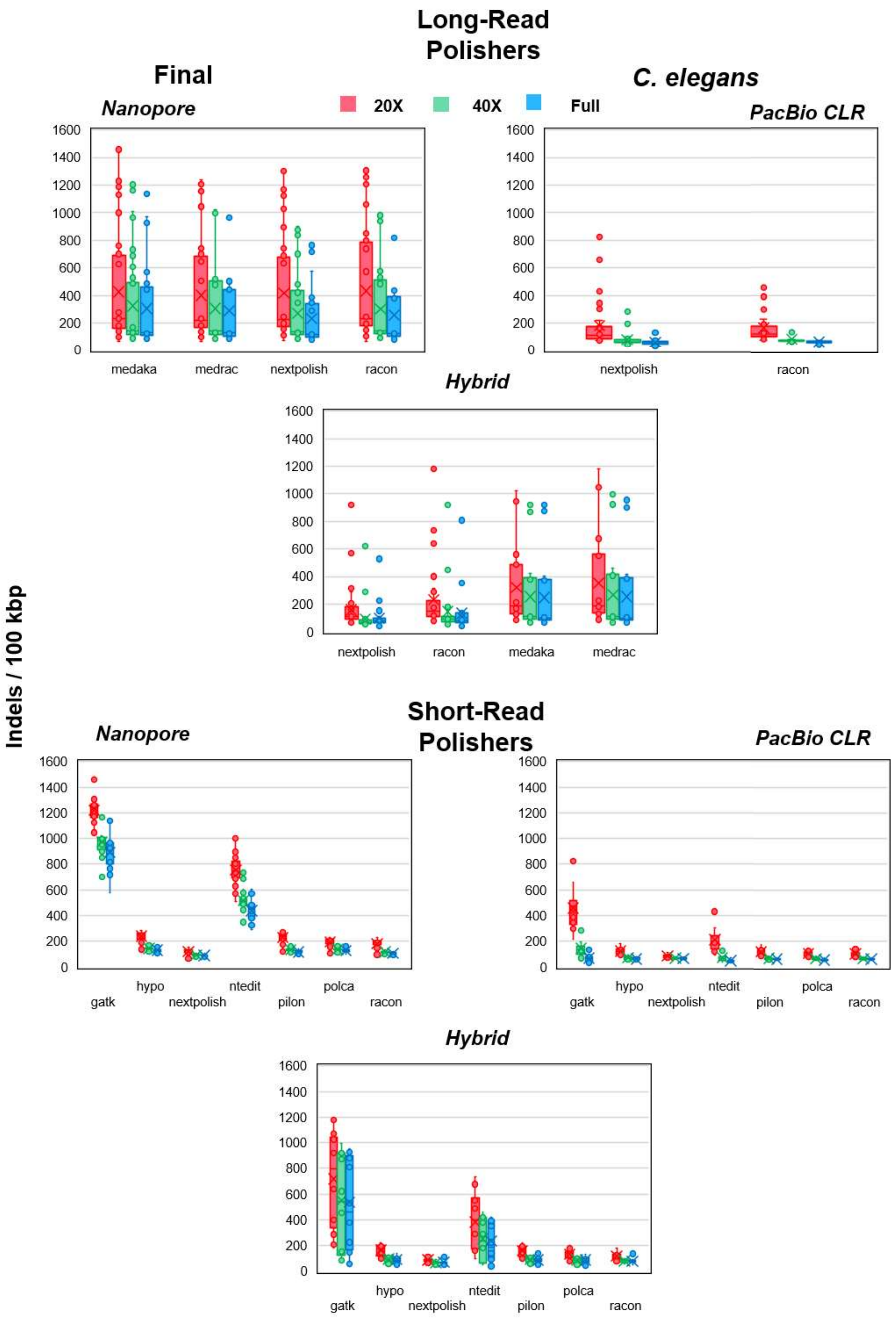


**Supplementary Figure S9:** Box-and-whisker plots of indel error rates per 100 kbp of sequence for all final assemblies for *C. elegans* at 20X (red), 40X (green) and full (blue) levels of coverage, by polishing algorithm.

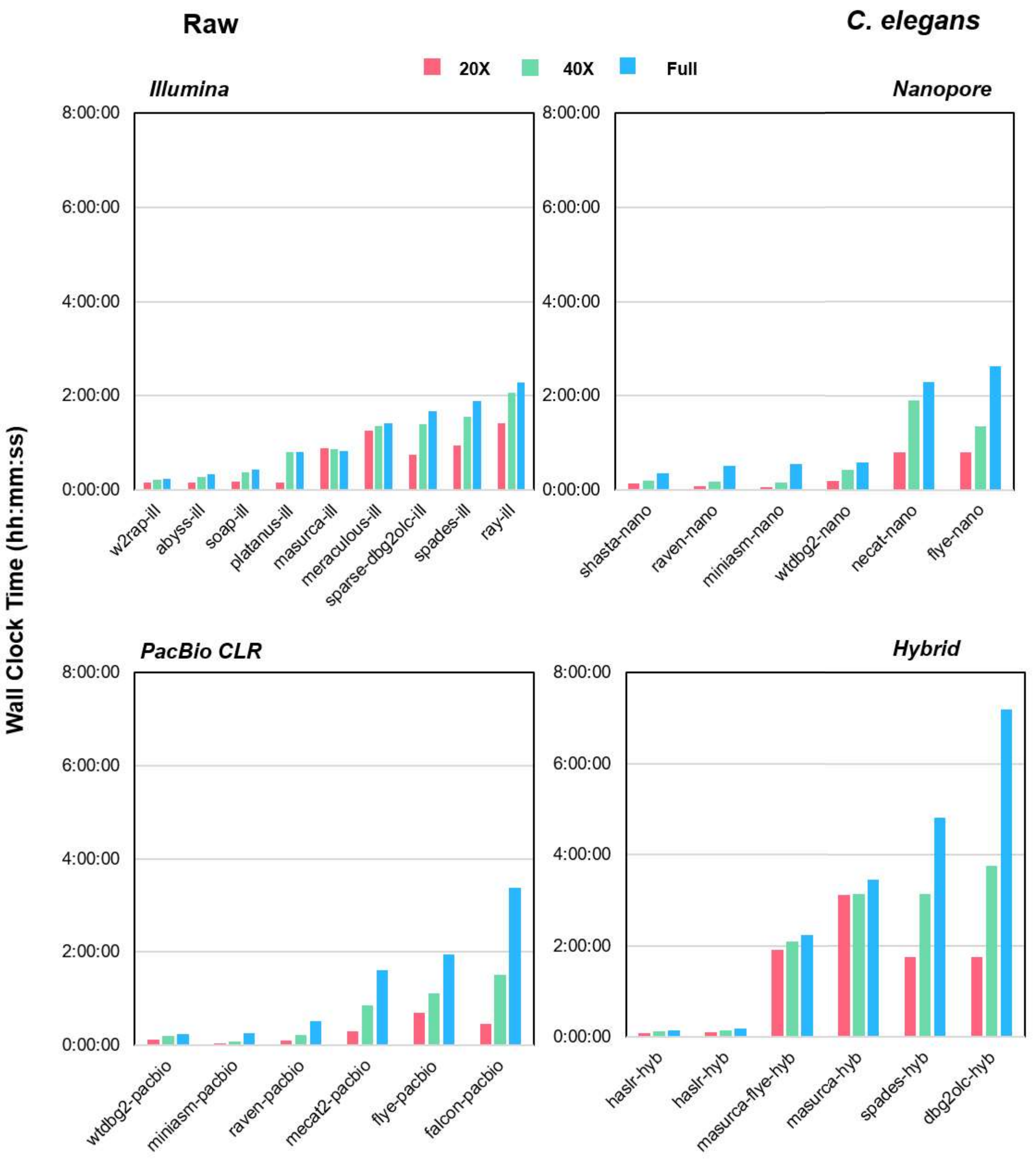


**Supplementary Figure S10:** Wall clock time of assembly algorithms used to produce raw assemblies for *C. elegans*. For Nanopore and PacBio CLR, times for Canu and Canu-Fast are not shown.

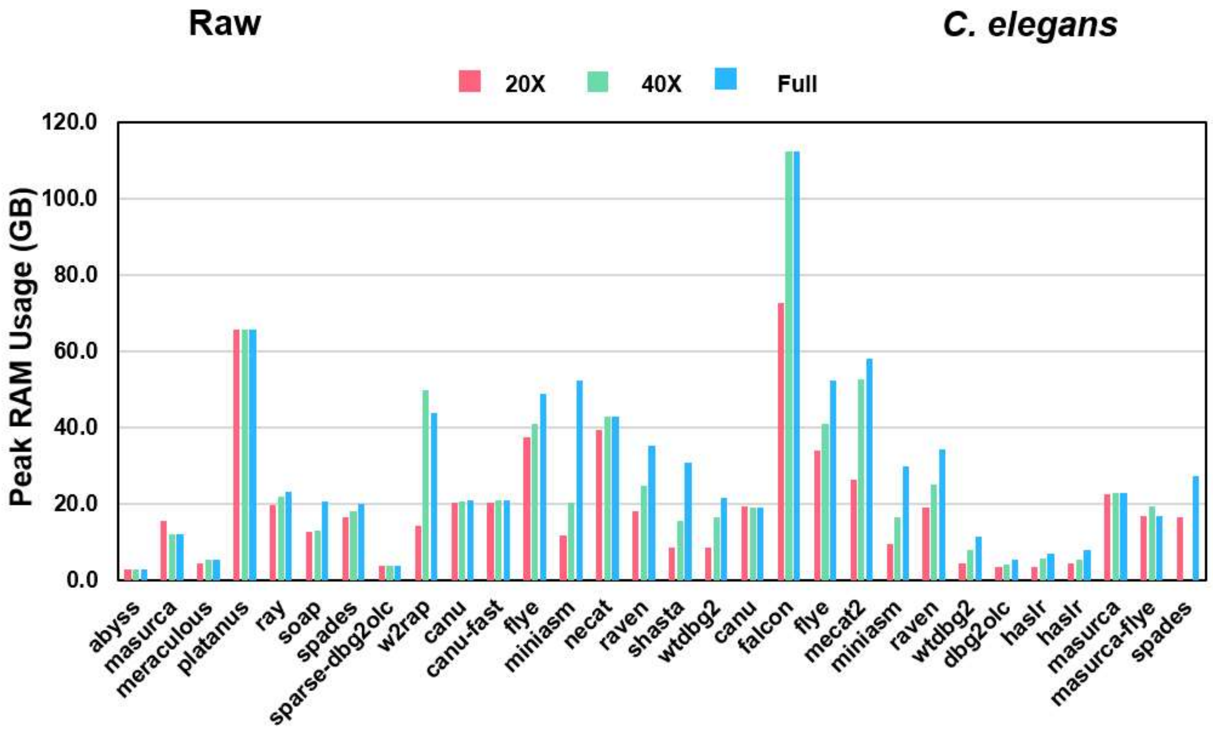


**Supplementary Figure S11:** Peak RAM usage of assembly algorithms used to produce raw assemblies for *C. elegans*.

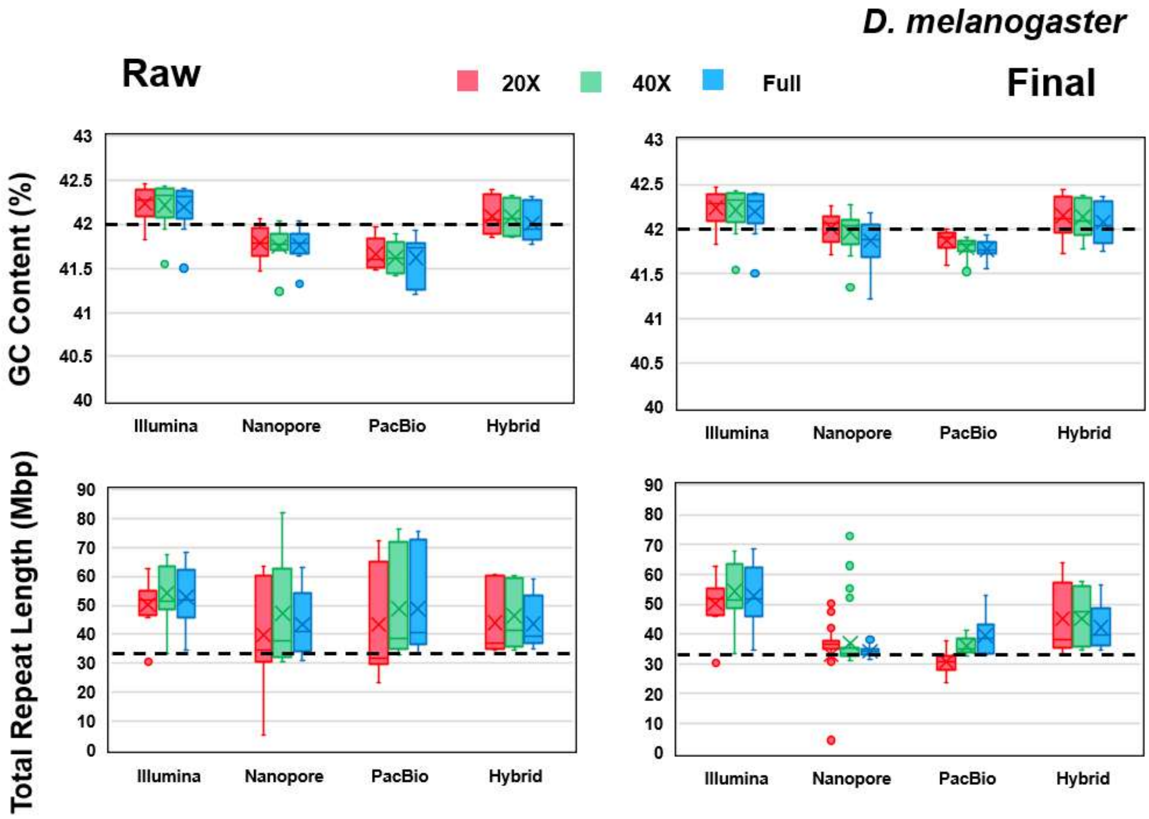


**Supplementary Figure S12:** Additional box-and-whisker plots of contiguity metrics for *D. melanogaster*, comparing GC% and total repeat length between raw and final assemblies. The dotted horizontal lines indicate the reference values.

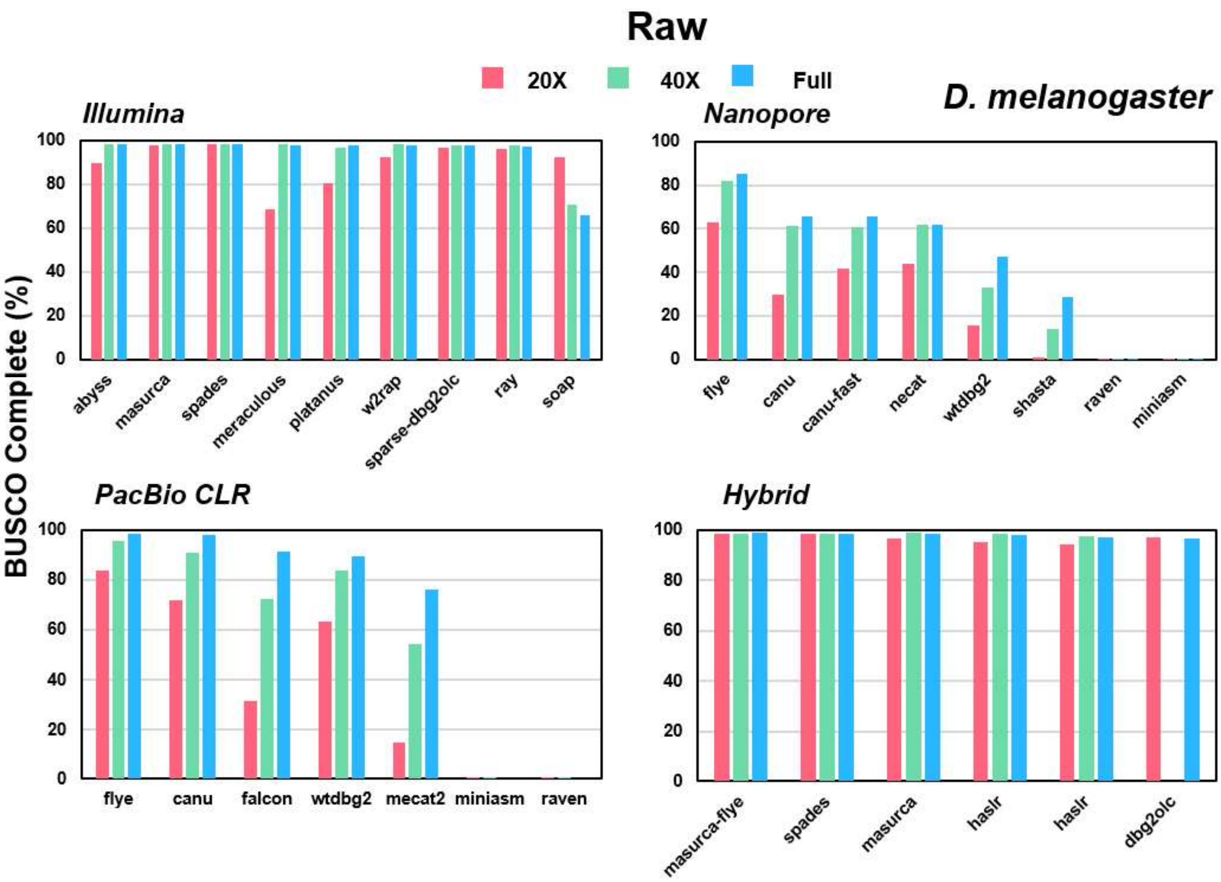


**Supplementary Figure S13:** BUSCO complete percentages for all raw assemblies for *D. melanogaster* at 20X (red), 40X (green) and full (blue) levels of coverage.

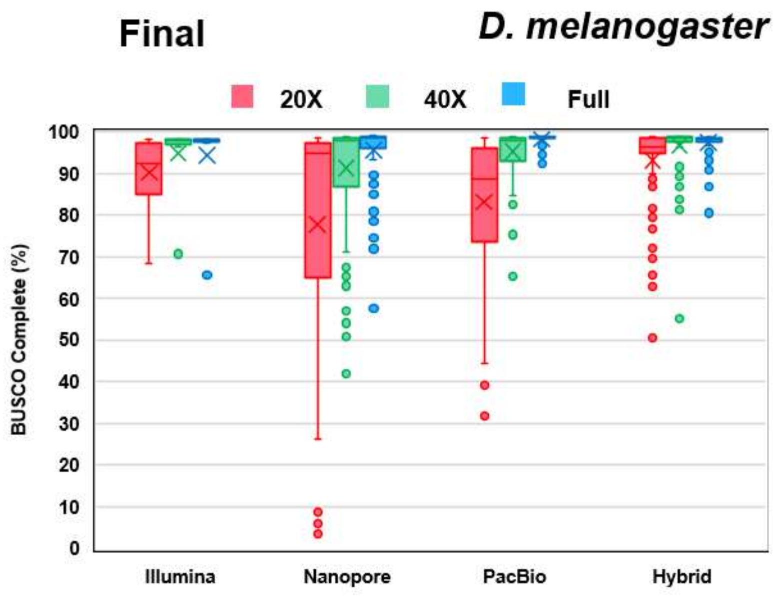


**Supplementary Figure S14:** Box-and-whisker plots of BUSCO complete percentages for all final assemblies for *D. melanogaster* at 20X (red), 40X (green) and full (blue) levels of coverage, by sequencing type.

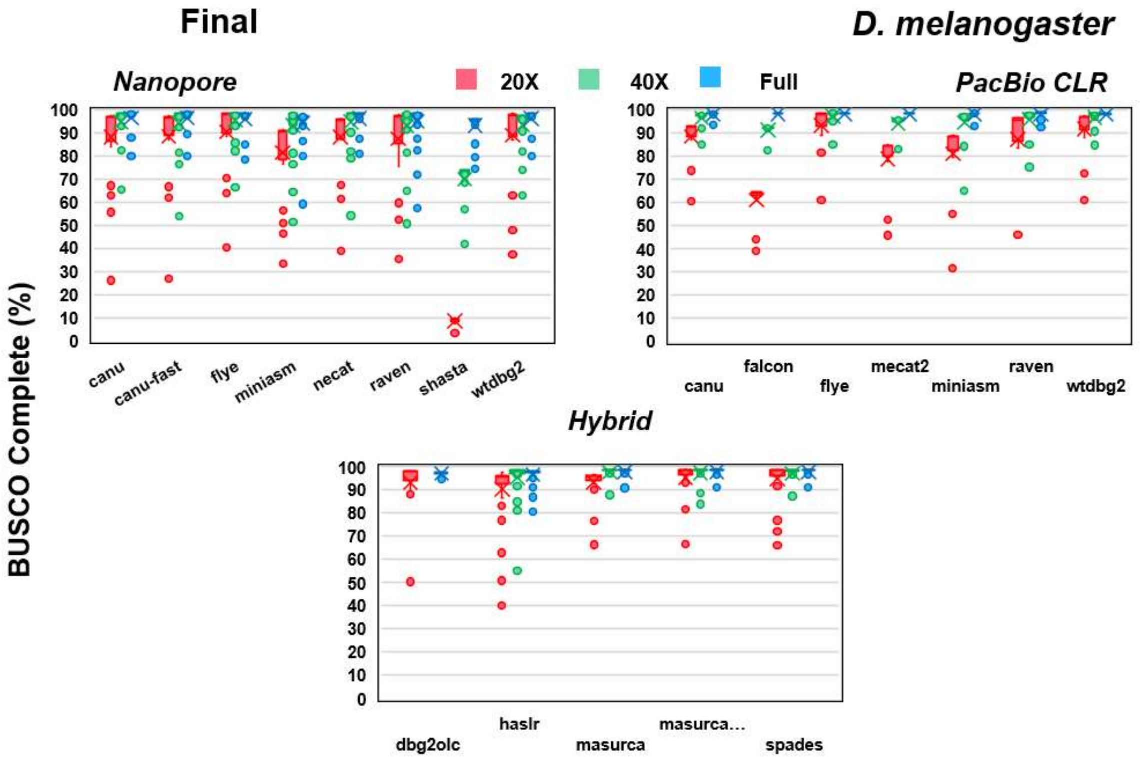


**Supplementary Figure S15:** Box-and-whisker plots of BUSCO complete percentages for all final assemblies for *D.melanogaster* at 20X (red), 40X (green) and full (blue) levels of coverage, by assembler.

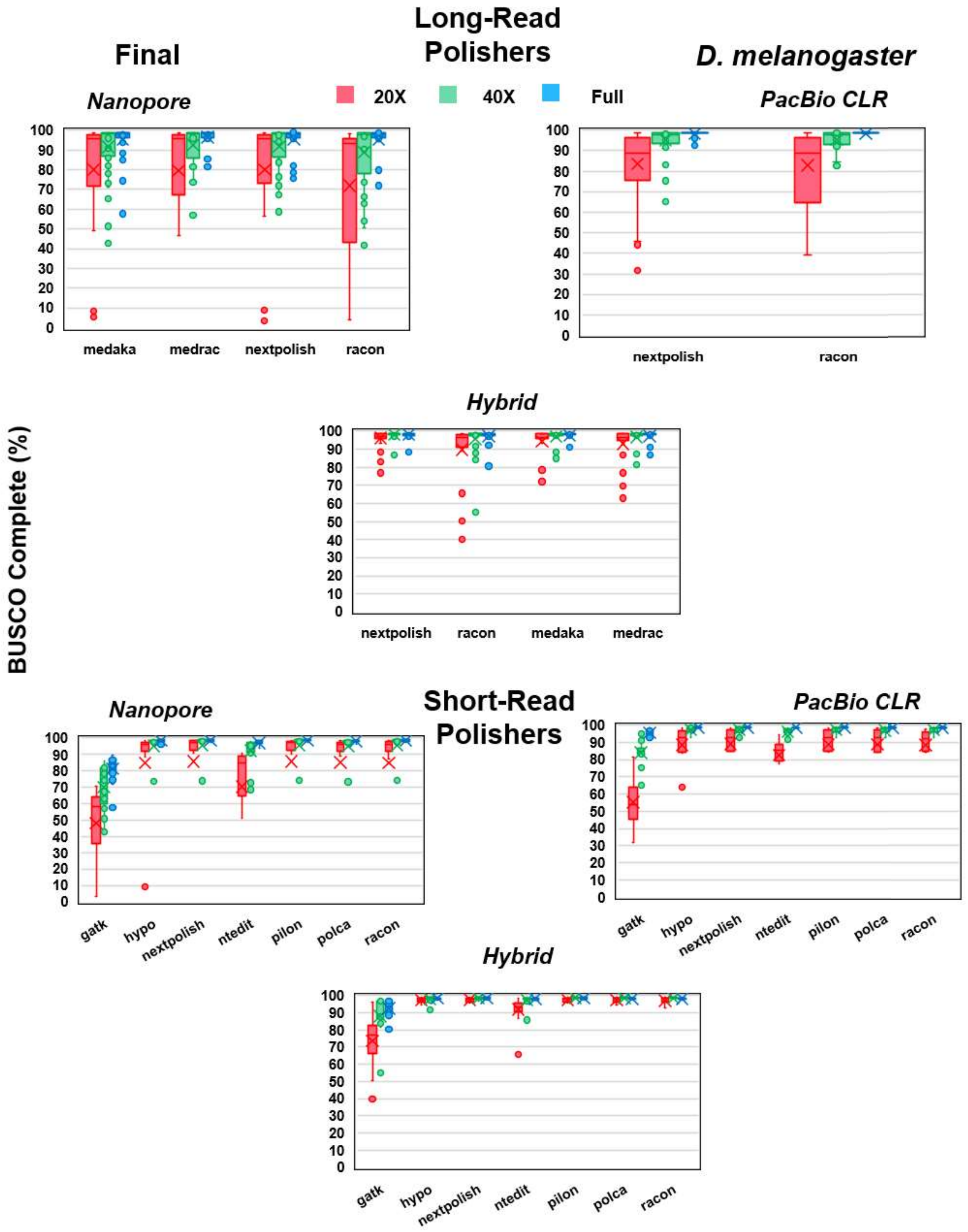


**Supplementary Figure S16:** Box-and-whisker plots of BUSCO complete percentages for all final assemblies for *D.melanogaster* at 20X (red), 40X (green) and full (blue) levels of coverage, by polishing algorithm used.

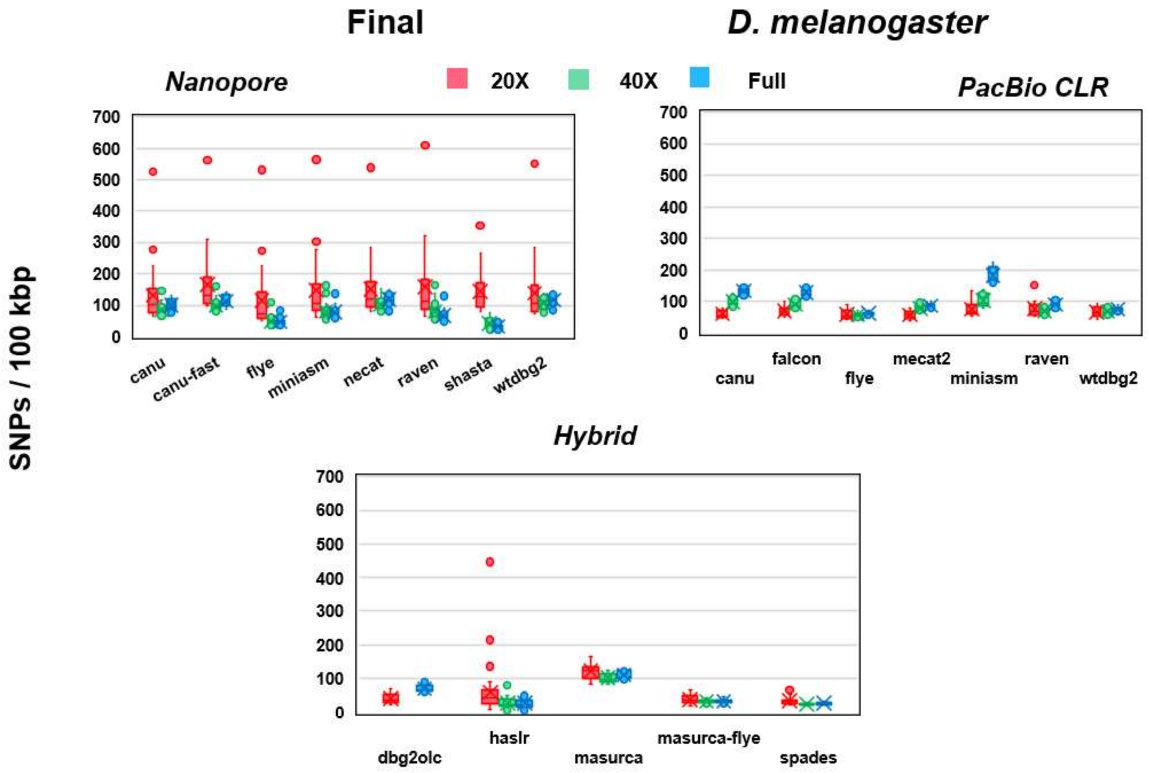


**Supplementary Figure S17:** Box-and-whisker plots of SNP error rates per 100 kbp of sequence for all final assemblies for *D. melanogaster* at 20X (red), 40X (green) and full (blue) levels of coverage, by assembler.

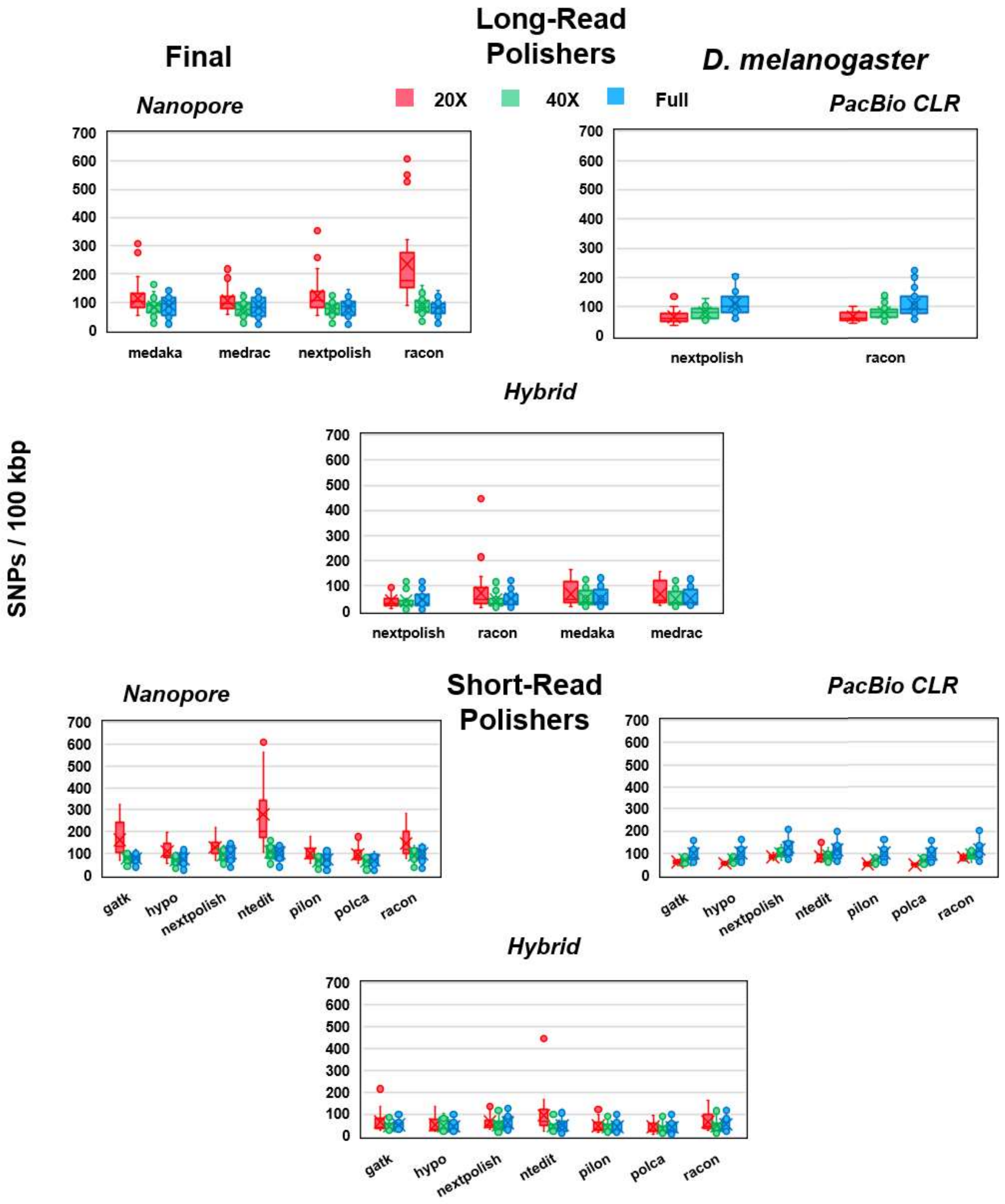


**Supplementary Figure S18:** Box-and-whisker plots of SNP error rates per 100 kbp of sequence for all final assemblies for *D. melanogaster* at 20X (red), 40X (green) and full (blue) levels of coverage, by polishing algorithm used.

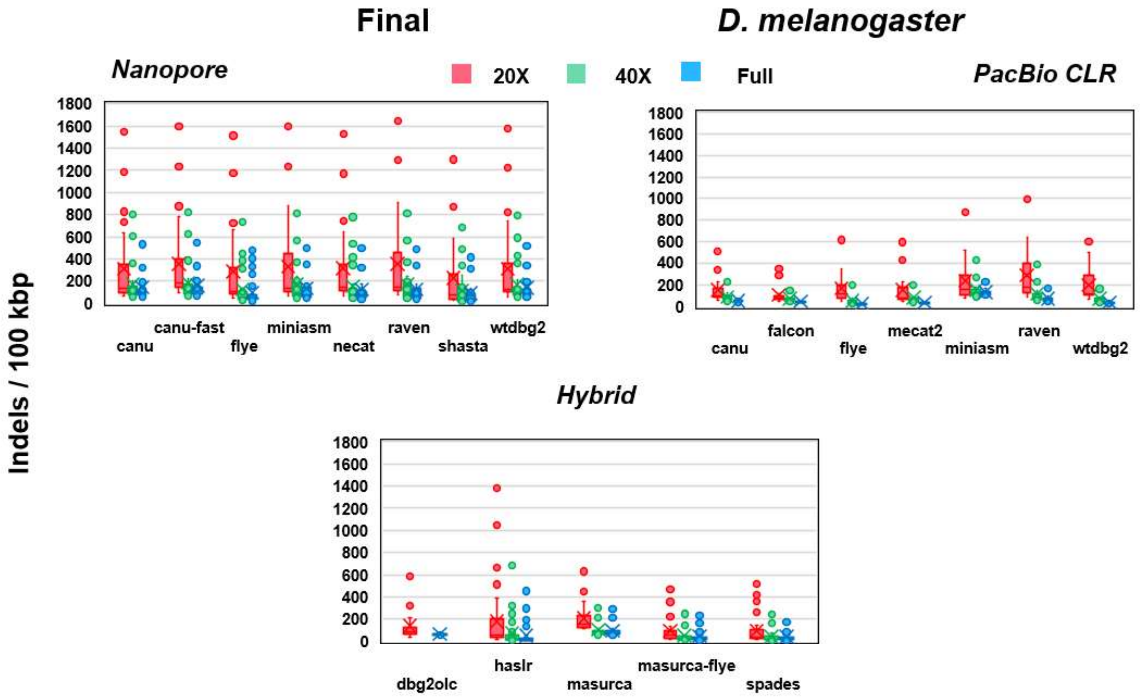


**Supplementary Figure S19:** Box-and-whisker plots of indel error rates per 100 kbp of sequence for all final assemblies for *D. melanogaster* at 20X (red), 40X (green) and full (blue) levels of coverage, by assembler.

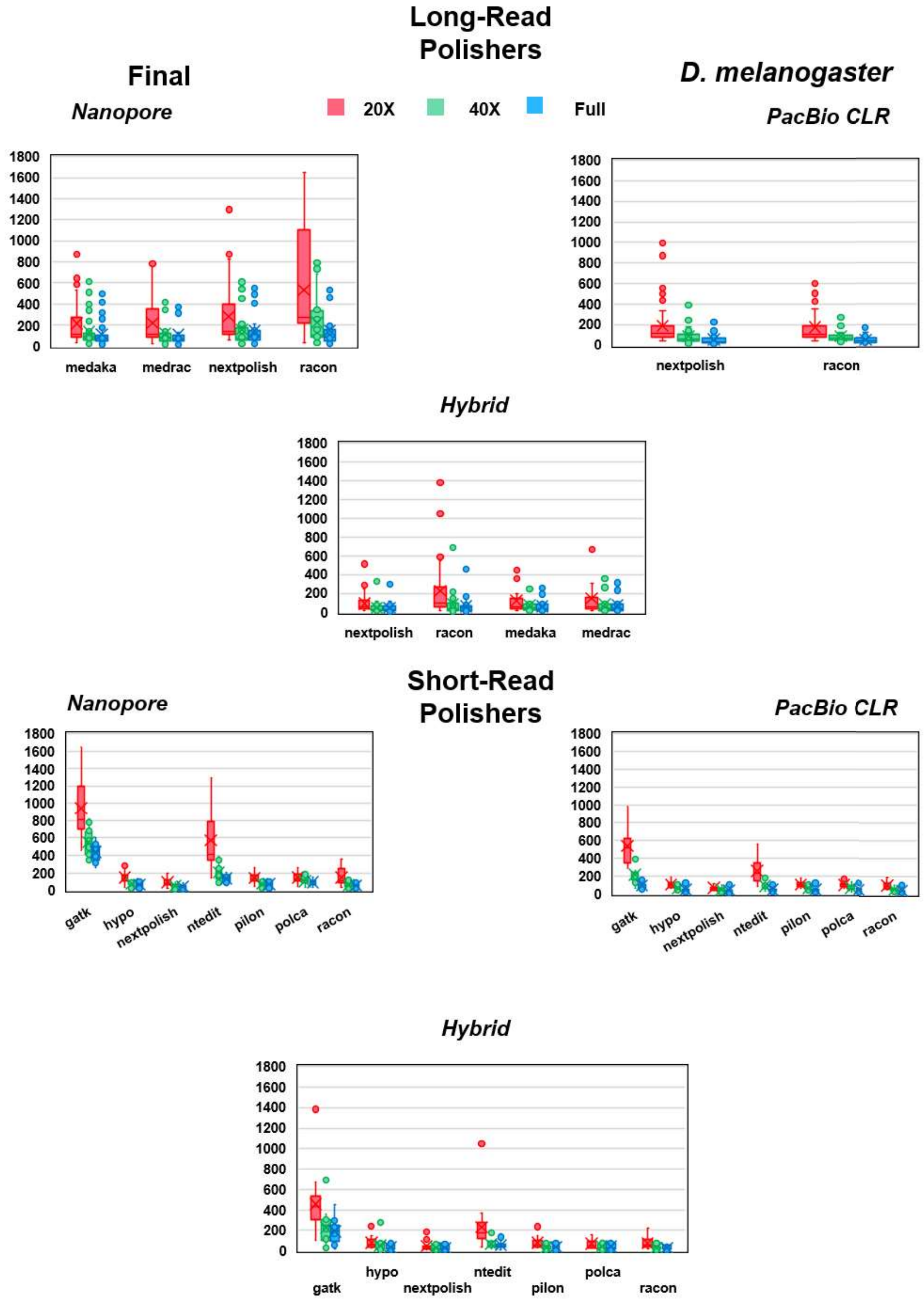


**Supplementary Figure S20:** Box-and-whisker plots of indel error rates per 100 kbp of sequence for all final assemblies for *D. melanogaster* at 20X (red), 40X (green) and full (blue) levels of coverage, by polishing algorithm used.

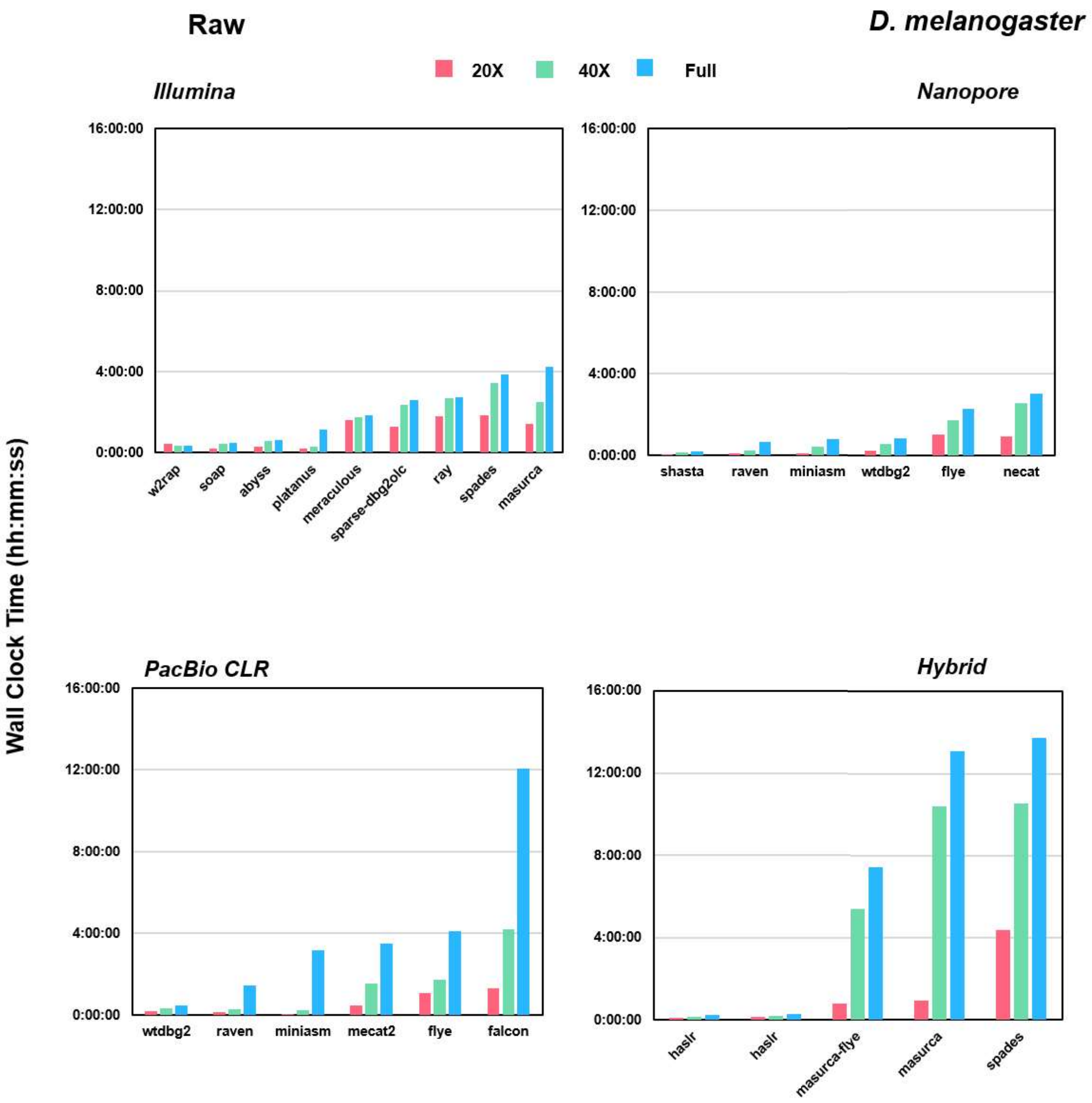


**Supplementary Figure S21:** Wall clock time of assembly algorithms used to produce raw assemblies for *D. melanogaster*. For Nanopore and PacBio CLR, times for Canu and Canu-Fast are not shown, and for Hybrid, times for DBG2OLC are not shown.

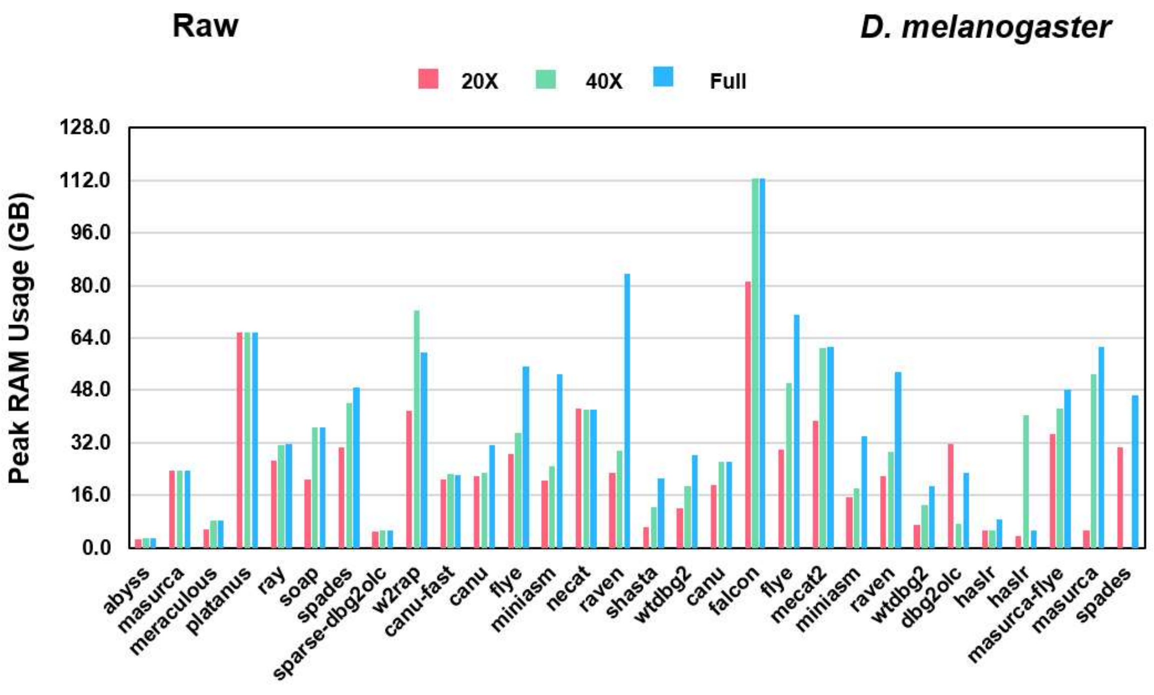


**Supplementary Figure S22:** Peak RAM usage of assembly algorithms used to produce raw assemblies for *D. melanogaster*.

#### 2. Supplementary Materials

##### Data Preparation

For Illumina data, reads were trimmed and quality-filtered with trim-galore v 0.6.4 using the following parameters:

trim_galore --cores 4 --quality 20 --length 90 --trim-n --max_n 0 --paired {reads}

For Nanopore and PacBio CLR data, reads were quality filtered with Filtlong v 0.2.0 using the following parameters:

filtlong --min_length 1000 --keep_percent 90 {reads}

---

##### Assembly

##### Illumina Assemblies

The following parameters were used for each assembler benchmarked in this study, with the same parameters for *C. elegans* and *D. melanogaster* unless noted:

**ABySS**:

abyss-pe name={prefix} k=96 in='{reads}' j={threads} B=2G H=4 kc=2 v=-v

**MaSuRCA**:

masurca {config};
bash assemble.sh;
[uses config file – please see https://github.com/genomeassembler/benchmarking-study for default]

**Meraculous**

run_meraculous.sh -c {config} -dir {workdir} -cleanup_level=1
[uses config file – please see https://github.com/genomeassembler/benchmarking-study for default]

**Platanus**

platanus assemble -o {prefix} -f {reads} -k 32 -t {threads} -m {memory};
platanus scaffold -o {prefix} -c {contigs} -b {bubbles} -IP1 {reads} -t {threads};
platanus gap_close -o {prefix} -c {scaffolds} -f {reads} -t {threads};

**Ray**

mpiexec -n {threads} Ray -p {reads} -o {outdir}

**SOAPdenovo2**

SOAPdenovo-63mer all -s {config} -o {outdir} -K 51 -p {threads} -N {genome-size}
[uses config file – please see https://github.com/genomeassembler/benchmarking-study for default]

**SPAdes**

spades.py -1 {reads} -2 {reads} -t {threads} -m {memory} -o {outdir}

**SparseAssembler + DBG2OLC**

SparseAssembler g 10 k 51 LD 0 GS {genome-size} f {reads} f {reads};
DBG2OLC LD1 0 Contigs {contigs} k 31 KmerCovTh 0 MinOverlap 50 PathCovTh 1 f {reads} f {reads};

**w2rap**

w2rap-contigger -t {threads} -m {memory} -o {outdir} -r {reads} -p {prefix}

---

**Long-Read Assemblies**

The following parameters were used for each assembler benchmarked in this study, with the same parameters for *C. elegans* and *D. melanogaster* unless noted:

**Canu**

canu -p {prefix} -d {outdir} genomeSize={genome-size} –{pacbio or nanopore}-raw {reads} -useGrid=false;

**Canu (fast mode)**

canu -p {prefix} -d {outdir} genomeSize={genome-size} –{pacbio or nanopore}-raw {reads} -useGrid=false -fast;

**FALCON**

fc_run falcon.cfg
[uses config file – please see https://github.com/genomeassembler/benchmarking-study for default]**Flye**

flye --{nanopore or pacbio}-raw {reads} --out-dir {outdir} --threads {threads}

**MECAT2**

mecat.pl correct {config};
mecat.pl trim {config};
mecat.pl assemble {config};
[uses config file – please see https://github.com/genomeassembler/benchmarking-study for default]

**Miniasm**

minimap2 -x ava-{ont or pb} -t {threads} {reads} {reads};
miniasm -f {reads} {align};

**NECAT**

necat.pl bridge {config};
[uses config file – please see https://github.com/genomeassembler/benchmarking-study for default]

**Raven**

raven -p 0 -t {threads} {reads}

**Shasta**

shasta --input {reads} --threads {threads}

**WTDBG2**

wtdbg2 -x {ont or rs} -g {genome-size} -t {threads} -i {reads} -fo {prefix};
wtpoa-cns -t {threads} -i {contigs} -fo {outfile};

---

**Hybrid Assemblies**

The following parameters were used for each assembler benchmarked in this study, with the same parameters for *C. elegans* and *D. melanogaster* unless noted:

**DBG2OLC**

SparseAssembler g 10 k 51 LD 0 GS {genome-size} f {reads} f {reads};
DBG2OLC Contigs {contigs} k 17 KmerCovTh 2 MinOverlap 50 AdaptiveTh 0.01
RemoveChimera 1 f {reads};
cat {contigs} {reads} > {all};
split_reads_by_backbone.py -b {backbone} -o {workdir} -r {all} -c {consensus-info};
split_and_run_sparc.sh {backbone} {consensus-info} {all} {workdir} 2 > {log};

**HASLR**

python haslr.py -t {threads} -o {outdir} -g {genome-size} -l {longreads} -x {nanopore or pacbio} -s {shortreads}

**MaSuRCA**

masurca {config};
bash assemble.sh;
[uses config file – please see https://github.com/genomeassembler/benchmarking-study for default]

**MaSuRCA + Flye**

masurca {config};
bash assemble.sh;
[uses config file – please see https://github.com/genomeassembler/benchmarking-study for default]

**SPAdes**

spades.py -1 {reads} -2 {reads} --nanopore {nanoreads} --pacbio {pacbioreads} -t {threads} -m {memory} -o {outdir}

---

**Polishing Packages**

The following parameters were used for each polishing algorithm benchmarked in this study, with the same parameters for *C. elegans* and *D. melanogaster* unless noted:

**Short-Read Polishing**

**GATK**

bwa index {assembly.fa};

bwa mem -t {threads} {assembly.fa} {reads} | samtools sort -o {bamfile} -@ {threads} -;

samtools index {bamfile};

bash ../../../../scripts/gatk-variant.sh asm.fa out1.bam out1.vcf;

picard CreateSequenceDictionary R= {assembly.fa} O= {assembly.dict};

samtools faidx {assembly.fa}

gatk IndexFeatureFile -F {vcf-file}

gatk FastaAlternateReferenceMaker -O {polished.fa} -R {assembly.fa} -V{vcf-file}

**HyPo**

minimap2 --secondary=no --MD -ax sr -t {threads} {assembly.fa} {reads} | samtools view -Sb - | samtools sort -o {bamfile} -@ {threads} -;

samtools index {bamfile};

hypo -d {assembly.fa} -r @input.fofn -s {genomesize} -c {readcoverage} -b {bamfile} -p 80 -t {threads} -o {polished.fa};

**NextPolish**

echo -e "{reads}" > sgs.fofn;

echo -e "job_type = local\njob_prefix = run-1\ntask = 12\nrewrite = yes\nmultithread_jobs = 5\nparallel_jobs = 4\ngenome = ./{assembly.fa}\ngenome_size = auto\nworkdir = ./01_rundir\npolish_options = -p {{multithread_jobs}}\nsgs_fofn = sgs.fofn\nsgs_options = -bwa" > run-1.cfg;

nextPolish run-1.cfg;

**ntEdit**

nthits -b 36 -k 50 -t {threads} -p ntedit-bloom --outbloom --solid @inputs.fofn;

ntedit -k 50 -f {assembly.fa} -r ntedit-bloom*.bf -b {polished} -t {threads};

**Pilon**

bwa index {assembly.fa};

bwa mem -t {threads} {assembly.fa} {reads} | samtools sort -o {bamfile} -@ {threads} -;

samtools index {bamfile};

java -Xmx{memory}G -jar pilon.jar --genome {assembly.fa} --frags {bamfile} --output {polished} --fix all --threads {threads};

**POLCA**

bash MaSuRCA-3.3.7/bin/polca.sh -a {assembly.fa} -r '{reads}' -t {threads};

**Racon**

minimap2 -x sr {assembly.fa} {reads} > {paf-file}

racon -u -t {threads}{reads} {paf-file} {assembly.fa} > {polished.fa};

---

**Long-Read Polishing**

**Medaka**

medaka_consensus -i {reads} -d {assembly.fa} -o {out-dir} -t {threads};

**Racon + Medaka (‘medrac’)**

[Racon run as below for 4 iterations, followed by the medaka step above]

**NextPolish**

echo -e “{reads}" > lgs.fofn;

echo -e "job_type = local\njob_prefix = run-1\ntask = 5\nrewrite = yes\nmultithread_jobs = 5\nparallel_jobs = 4\ngenome = ./{assembly.fa}\ngenome_size = auto\nworkdir = ./01_rundir\npolish_options = -p {{multithread_jobs}}\nlgs_fofn = lgs.fofn\nlgs_minimap2_options = -x map-{ont or pb}" > run-1.cfg;

nextPolish run-1.cfg;

**Racon**

minimap2 -x map-{ont or pb} {assembly.fa} {reads} > {paf-file};

racon -u -t 20 {reads} {paf-file} {assembly.fa} > {polished.fa};

---

**Hybrid Polishing**

**HyPo**

echo -e '{short-reads}' > input.fofn;

minimap2 --secondary=no --MD -ax sr -t {threads} {assembly.fa} {short-reads} | samtools view -Sb - | samtools sort -o {short-read-bam-file} -@ {threads} -;

samtools index {short-read-bam-file}; ';

minimap2 --secondary=no --MD -ax map-pb -t {threads} {assembly.fa} {long-reads} | samtools view -Sb - | samtools sort -o {long-read-bam-file} -@ {threads} -;

samtools index {long-read-bam-file};

hypo -d {assembly.fa} -r @input.fofn -s {genomesize} -c {short-read-coverage} -b {short-read-bam-file}-B {long-read-bam-file}-p 80 -t {threads} -o {polished.fa};

---

**Metric Evaluation**

**assembly-stats**

assembly-stats {assembly.fa} > {assembly-stats-output.txt};

**BUSCO**

busco -i {assembly.fa} -o {assembly} -l {nematoda_odb9 or diptera_odb9} -m geno -c {threads};

**BUSCOMP**

[BUSCO run on each assembly above, with FASTA files for each assembly in fasta directory]

echo -e
'genomesize={genomesize}\nruns="run_*,../ref/run_ref"\nfastadir=fasta/\nbasefile=buscomp-out \nrmdreport=F\nfullreport=F\nforks={threads}' > input.ini

python buscomp.py -ini input.ini i=-1 ;

**QUAST**

python quast.py -o {outdir} -r {reference} -t {threads} -m 200 --extensive-mis-size 7000 --min-contig 3000 --min-alignment 500 --space-efficient --eukaryote --fast -l {labels} -1 {short-reads} -2 {short-reads} --{pacbio or nanopore} {long-reads} {assemblies.fa}

**Red**

Red -gnm ./ -msk ./masked-Red -cor {threads} 1> {output.txt};
